# MMCCI: multimodal integrative analysis of single-cell and spatial cell-type communications to uncover overarching and condition-specific ligand-receptor interaction pathways

**DOI:** 10.1101/2024.02.28.582639

**Authors:** Levi Hockey, Onkar Mulay, Zherui Xiong, Samuel X. Tan, Kiarash Khosrotehrani, Christian M. Nefzger, Quan Nguyen

## Abstract

Cell-cell interaction (CCI) analyses are an indispensable tool for harnessing the detail and depth of spatial and single-cell transcriptomics datasets by inferring inter-cellular communications, but no methods to integrate CCI results across samples exist currently. To address this, we have developed a computational pipeline, Multimodal CCI (MMCCI), to statistically integrate and analyze CCI results from existing popular CCI tools. We benchmarked MMCCI’s integration on single-cell spatial datasets and found it to be highly accurate compared to simpler methods. We utilized MMCCI’s integration and downstream biological analyses to uncover global and differential interaction patterns in multimodal aging brain and melanoma spatial datasets.

## Background

Spatial transcriptomics (ST) is a molecular profiling technique that maps RNA sequencing with spatial information to access spatial gene expression patterns. This information can provide deep biological insights into many applications including neuroscience and cancer research [1, 2]. A valuable method to utilise ST data is cell-cell interaction (CCI) analysis, which aims to decipher the intricate communication networks within tissues and understand how cells interact in their spatial context. Multiple computational tools exist for inferring and analyzing intercellular communication networks in single-cell RNA-sequencing (scRNA-seq) and ST data such as stLearn, CellChat, Squidpy, CellPhoneDB, NATMI, and NicheNet [3, 4, 5, 6, 7, 8]. Tools such as stLearn and CellChat take spatial information into account, limiting the inferred communications to between only nearby cells or spots, whereas Squidpy, CellPhoneDB and NATMI infer communications between any given cells or spots and are thus prone to false discoveries, which is a major challenge in CCI analyses. Tools such as scDiffcom and COMUNET perform additional downstream analyses on interaction networks such as differential LR interactions and CCI. [9, 10].

For single cell data, integrative analysis of multiple modalities has shown to uncover complex and functionally important molecular and cellular mechanisms [11]. ST technologies such as CosMx and Xenium provide single cell resolution at the cost of only being able to detect a smaller panel of genes or proteins, while Visium and bin80 STOmics (bin80 produces equivalent size to Visium spot at 55um wide) are able to sequence the entire transcriptome, but each Visium spot (55um wide) captures the expression of often one to ten cells [12, 13, 14, 15]. We posit that integration of these orthogonal technologies can therefore enable detection of a large number of spatially proximate LR pairs while still retaining the specificity of single cell spatial methods, providing deep insights into biological processes that are otherwise unachievable from any one technology alone. However, qualitative or ‘average-based’ analyses are vulnerable to false positives and single-sample bias, and there are currently no methods to integrate CCI results from multiple samples across shared or different transcriptomic modalities.

Therefore, we introduce MMCCI as a statistical framework for multimodal CCI integration. MMCCI is based on the assumption that the integration of CCI results across different spatial transcriptomics technologies or multiple samples will be able to filter false positive interactions while preserving the true interactions, thereby constructing a more accurate landscape of the complex intercellular communications than any one sample or modality alone. As the first pipeline for quantitatively integrating CCI results across transcriptomic platforms and from multiple samples, MMCCI is able to provide highly confident CCI results through statistical meta-analysis, combining p-values for each interaction across each sample, as well as batch-effect correction, adjusting for unwanted technical differences between samples both within and across modalities. MMCCI provides a wide range of novel functionalities, including:

- First tool for statistical integration and meta-analysis of multimodal spatial and single-cell transcriptomics CCI results from existing popular CCI tools.
- Comparative analysis of integrated or individual cell-type networks between different groups to identify the differences in cell-cell LR interactions between biological conditions.
- Novel tools for delving into specific interactions and the biological pathways involved through enrichment analysis, including cell-type pair LR querying and clustering of LRs based on cell-type networks to help infer clear biological patterns from complex CCI results.
- Analyzing cell types and LR pairs involved in specific biological pathways enriched in CCI results, thereby enabling hypothesis testing about the roles of cells and LRs involved in such pathways.
- Spatial clustering of cells/spots based on LR interaction scores to observe anatomical regions with similar interaction patterns, providing novel LR signatures for defining tissue regions.

## Results

### MMCCI integrates and analyses CCI results from multimodal datasets

MMCCI’s integration pipeline, shown in **Fig. 1a** aims to combine the interactions discovered in individual samples from existing popular CCI tools, such as stLearn, CellChat, CellPhoneDB, Squidpy, NATMI, and NicheNet, in order to identify global interaction patterns across multiple samples across multiple modalities. This integration pipeline can work across samples from either a single modality, or from samples across multiple modalities, such as combining scRNA-seq data and spatial transcriptomics data from different technologies like Visium and Xenium. MMCCI’s set of downstream analyses aim to provide clear biological insights from the deep and complex CCI results through Enrichr pathway analysis, differential CCI network analysis, cell-type LR interaction querying, and CCI network and interaction score clustering (**Fig. 1b**). **Fig. 1c** shows a simple example of how the cell type interactions for a single LR pair, HLA-B:KIR2DL3, across three simulated spatial breast cancer samples are integrated into a single network, capturing the overall cell type interaction pattern. MMCCI’s integration is able to do this for every LR pair across all samples. The integration and analysis pipelines are outlined in depth in the Methods section.

**Fig. 1.**
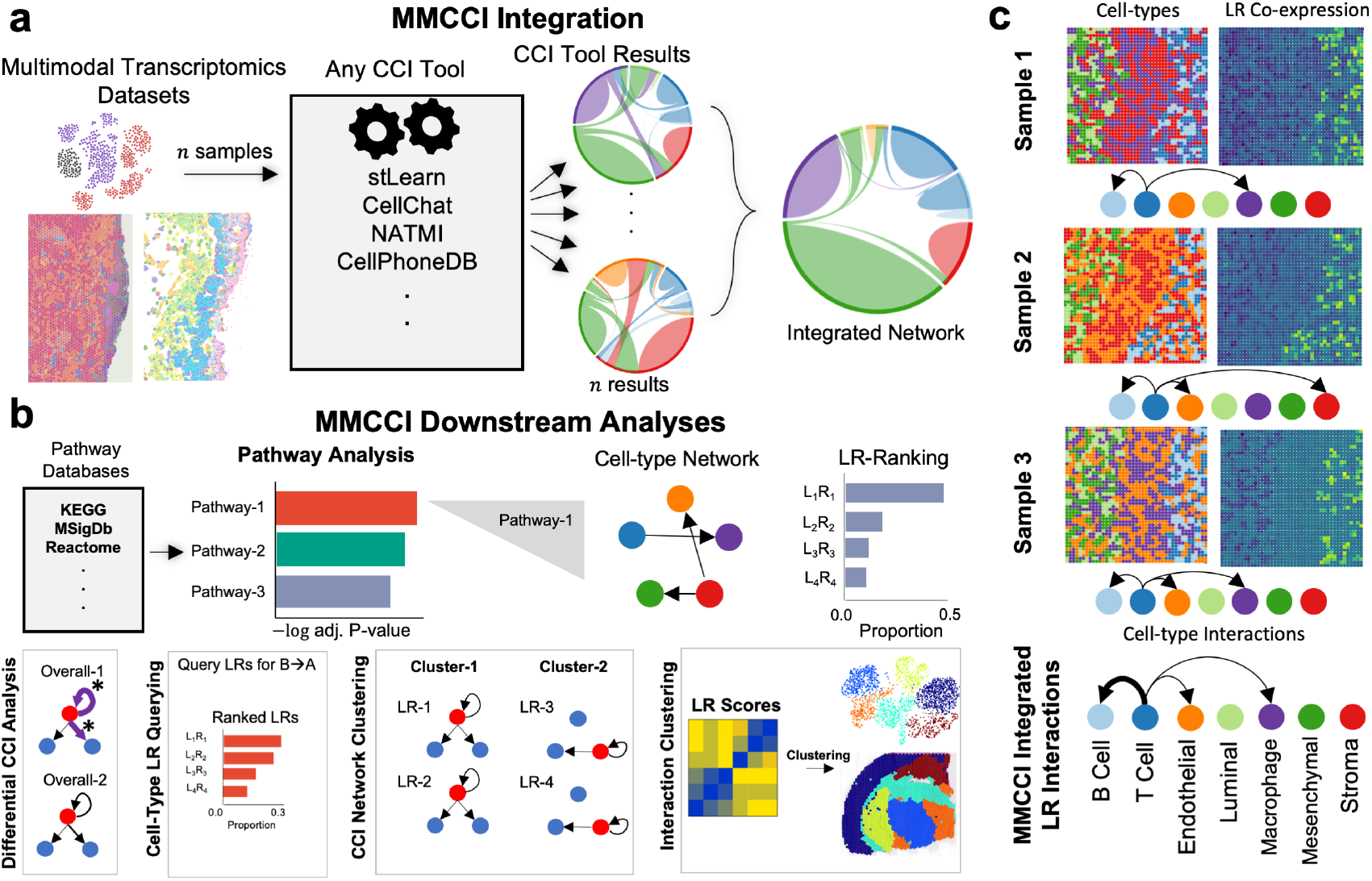
Overview of MMCCI’s integration method and analyses. **a**. Spatial or single-cell transcriptomics samples are individually processed through one of many different CCI tools. The CCI results from each sample is integrated into one CCI result through MMCCI integration. **b**. MMCCI’s downstream analyses. Shown at the top, Enrichr pathway analysis is run on the LR-pairs from the integrated result to identify significant biological pathways and the LR interactions involved. Shown from the bottom left to right, MMCCI also provides analysis functions including differential CCI analysis with statistical testing, LR pair querying and ranking for a sender-receiver cell-type pair, clustering of LR pairs with similar cell-type networks, and clustering of sample spots with similar interaction scores. **c**. An example of how MMCCI’s integration functions. Each spatial transcriptomics sample (simulated from scRNA-seq breast cancer) is run through a CCI algorithm (stLearn in this example). The integration of a single LR pair (HLA-B:KIR2DL3) is shown along with cell-type and LR co-expression spatial plots for each sample. The thicker arrows in the integrated result represents stronger interactions.

### Benchmarking and validation of MMCCI integration on single-cell whole-transcriptome spatial pancreas data

To validate the integration method, the latest and publicly available CosMx single-cell spatial, whole-transcriptome pancreas data consisting of 18 FOVs was used. The stLearn CCI results were compared to the MMCCI integration of the CCI results for each FOV across a number of quantitative and qualitative metrics (**Fig. 2a**) [12]. The CCI result of the entire sample was used as the “ground truth” as this contained the exact number of cell-type interactions for each LR pair across the whole sample as well as p-values for each interaction based on the background expression across the whole sample. The aim of the integration method was to match the integrated FOV CCI results to the whole sample CCI results as closely as possible, as this would show that the integration neither filters out important interactions nor retains false positive interactions which can be challenging when interaction counts and their p-values for specific LR pairs can vary greatly between samples when cell-type proportions and gene expression across samples vary. The heatmaps in **Fig. 2a**, which show the interaction counts from one cell type to another, are nearly identical between the whole and integrated, showing that MMCCI integration preserves the global LR interaction count proportions. **Fig. 2b** shows similar results, this time showing how the LR interaction counts per cell-type in the integrated (green line) are much closer to the whole (blue line) when compared to each individual FOV (dotted red lines) and the average across the FOVs (dark red line). **Fig. 2c** shows that the integrated results are significantly more similar to the whole at the individual LR level when compared to the average for each LR pair across all FOVs. The data followed a normal distribution according to SciPy’s “normaltest”, and a between-groups t-test indicated a p-value of < 0.0001 [16]. **Fig. 2d** shows how for two given sender-receiver cell types, ductal to ductal cells (a pair with a high number of interactions that are consistent across all the FOVs) and delta to beta cells (a pair with a low number of interactions with high variation across the FOVs), the top LR pairs and proportions are similar. **Sup. Fig. 1** shows these same plots, but for each FOV, showing how MMCCI is able to integrate these interactions across many FOVs with high accuracy even when there is a large amount of variation between samples. Overall, these results are able to show that MMCCI’s integration method is accurate when taking even a large number of samples with a high number of cells and genes.

**Fig. 2.**
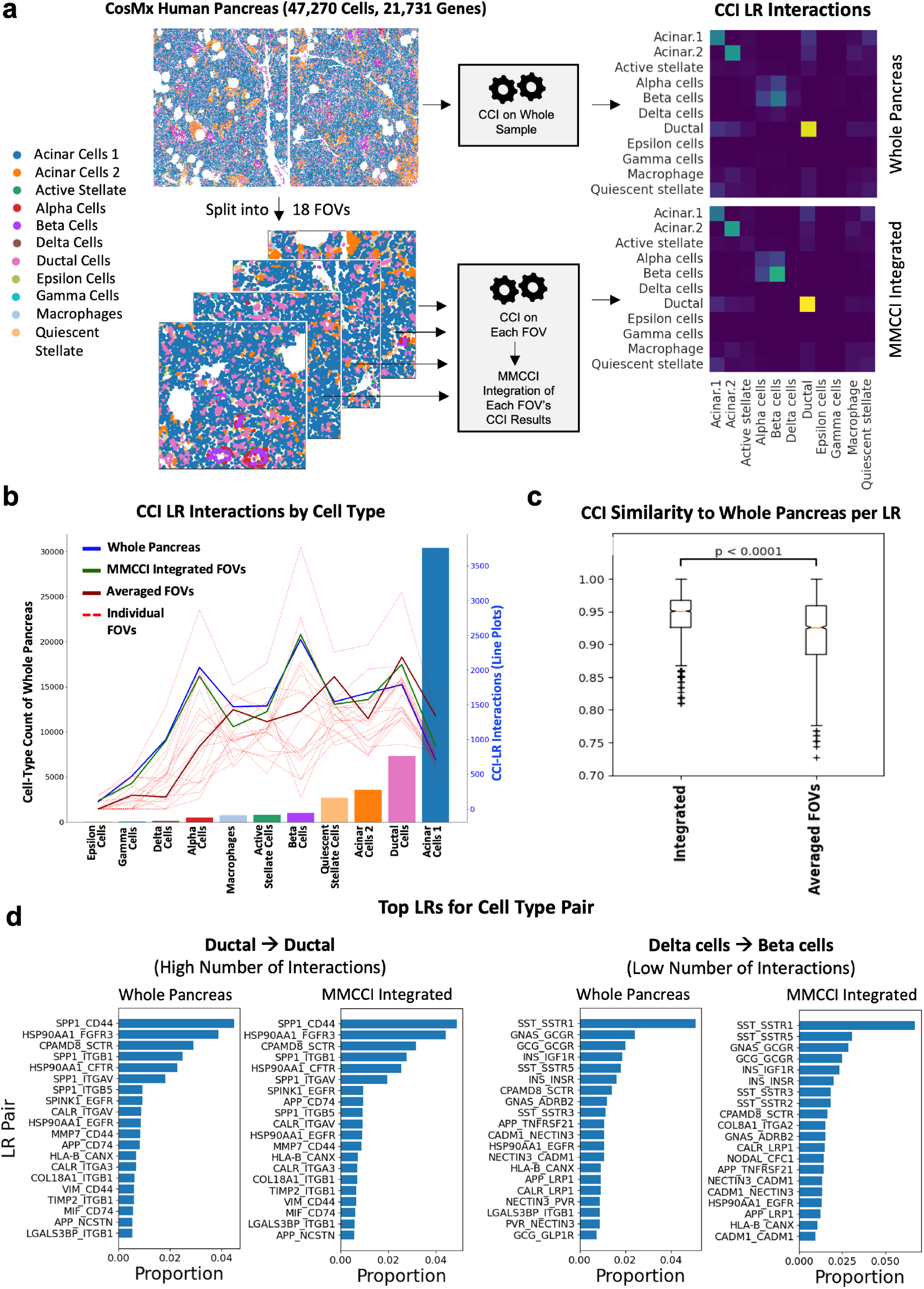
Benchmarking of MMCCI integration on CosMx human pancreas dataset. **a**. The entire sample was run through the CCI tool, stLearn, and compared to the MMCCI integration of each of the 18 FOVs run separately through stLearn. The heatmap of the global interaction counts for both the whole and integrated results are shown (right). **b**. Barplot of cell-type counts for the whole sample along with line plots showing the scaled CCI-LR interaction counts for each cell type for the whole pancreas (blue), MMCCI integrated FOVs (green), averaged FOVs (dark red), and individual FOVs (dotted red). **c**. The CCI matrix similarity scores per LR compared to the whole pancreas for MMCCI integrated and averaged across FOVs. **d**. Comparison of the top LR pairs and their proportions interacting in both the whole sample and MMCCI integrated result between ductal cells, which is a cell-type pair with a high number of interactions in the whole sample, and from delta to beta cells, which is a cell-type pair with a lower number of interactions in the whole sample.

### MMCCI integration and comparative analysis on multimodal spatial aging brain datasets

Next, MMCCI integration and downstream analyses were performed on the stLearn CCI results from our eight Visium mouse brain samples (four aged and four young) and four STOmics mouse brain samples (two aged and two young) (**Fig. 3**). These replicate samples were first integrated by technology and age group. The resulting four combined samples were again integrated by age group, resulting in one aged and one young mouse integrated network. Prior to integration, interactions between many cell types were observed interacting in individual samples, making it unclear which interactions were falsely identified and which were biological. After integration, only statistically significant interactions across the samples were retained (**Fig. 3**).

**Fig. 3.**
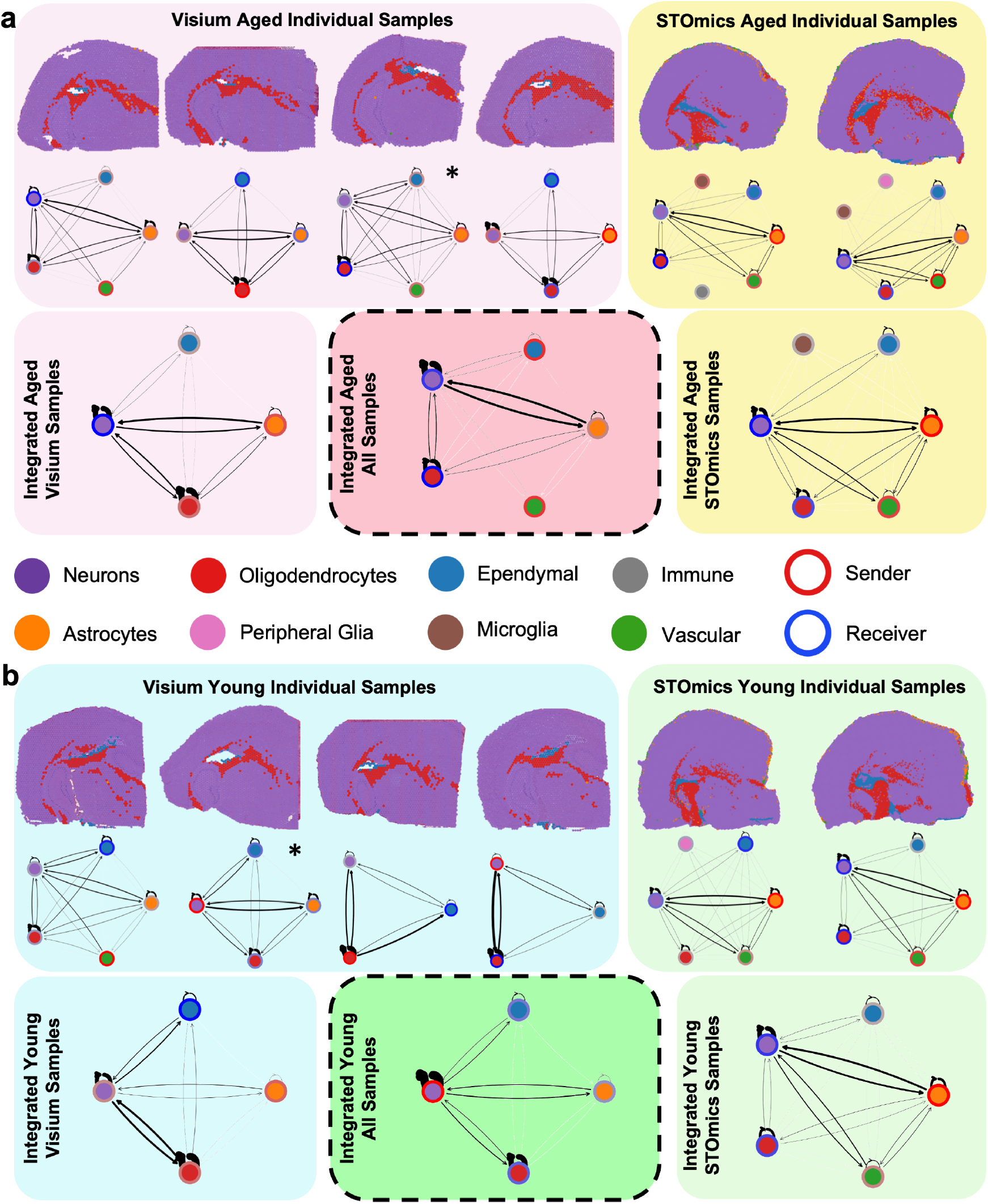
Integration of stLearn CCI results from aged and young mouse brain spatial transcriptomics samples. **a**. Cell-type spatial plots and overall stLearn interaction network plots of individual Visium (left) and STOmics (right) aged mouse brain results. The network with a (*) is the aged sample used in **Fig. 4f**. Below, the overall within-technology integration network of aged Visium (left) and aged STOmics (right) is shown along with the overall between-technology integration network of all aged samples (middle), which was used as the aged brain integrated sample for downstream analyses. In the network plots, the inner circle of the node shows the cell type and the outer ring shows if the cell type is sending more interactions (red) or receiving more interactions (blue) **b**. Similarly, young samples are shown.

#### MMCCI provides quantitative and qualitative differential interaction network analyses

A core feature of MMCCI is its pipeline for comparing cell-type interaction networks between different groups, highlighting how the roles of different cell-types and their interactions change in different biological conditions. For our aging dataset, the overall interaction networks for aged and young brain samples were constructed and the difference between these overall networks was calculated and run through MMCCI’s network permutation testing (see Methods section) to identify cell-type pair interactions that were significantly different between the aged and young samples, shown as the darker coloured edges in **Fig. 4a**. The lighter coloured edges show the differences that were not statistically significant (p-value *>* 0.05). The dissimilarity score between the overall aged and young networks was 0.232, which indicates that nearly a quarter of the network edges are significantly different between the age groups.

**Fig. 4.**
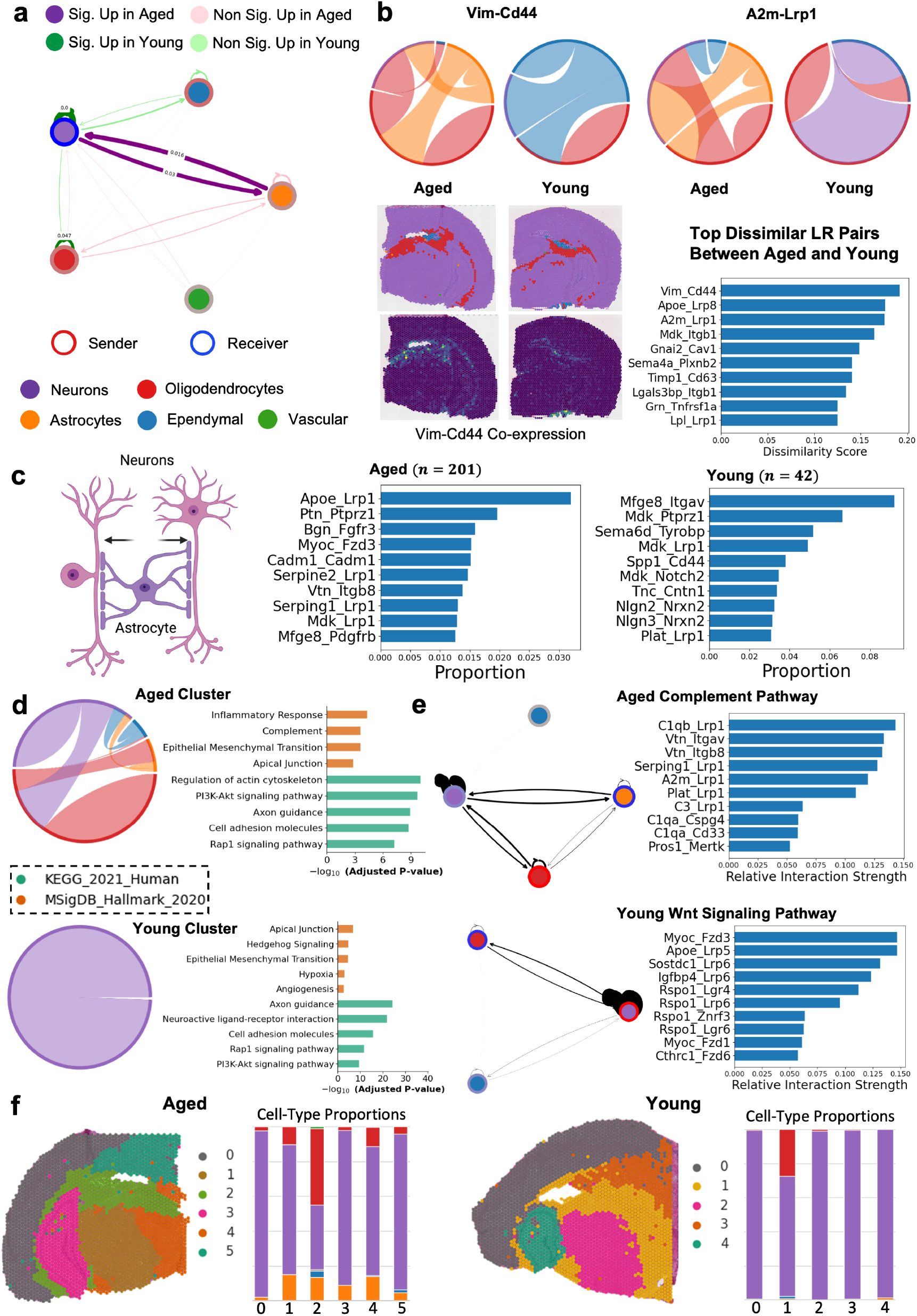
Downstream analysis of aged and young mouse brain CCI results. **a**. Overall network difference plot between aged and young, where significant edges are coloured and labeled with their p-value. The color of the edge indicates the age group that the interaction is increased in and the thickness of the edge indicates how much stronger the edge is upregulated in that age group. Edges with a p-value < 0.05 are darker in colour. **b**. Barplot of LR pairs (bottom right) with the most dissimilar cell-type interaction networks between aged and young with two LR pair’s networks for the integrated aged and young samples shown as chord plots. Chord plots are used to show the interacting cell types and the color of the chord indicates the cell type that is acting as mostly a sender. Spatial cell-type and Vim-Cd44 expression plots for an aged and a young Visium sample are shown. **c**. Barplot of top LR pairs interacting from astrocytes to neurons and their proportion of their interaction strength, shown for both aged and young. The number of LR pairs interacting from astrocytes to neurons in both aged groups is shown above the barplots. Created with BioRender.com. **d**. A pair of summed networks of clustered LR pairs from the integrated aged and young samples, where LR pairs with similar networks are clustered together, along with GSEA pathway analysis using Enrichr with the KEGG 2021 Human and MSigDB Hallmark 2020 databases. **e**. LR pairs with their summed networks involved in the complement pathway in the integrated aged sample and the Wnt signaling pathway in the young integrated sample. **f**. LR interaction score clustering results for an aged and a young Visium brain sample with bar plots showing the cell-type proportions of each cluster.

Interactions within ependymal cells were shown to decrease significantly with age, while interactions between astrocytes and neurons significantly increased with age. This reflects the current understanding that the ependymal layer thins during aging and reactive astrocytes proliferate and interpose themselves within the ependymal cell layer [17, 18]. Therefore, our findings are consistent with previous reports that interactions both within astrocytes and between astrocytes and neurons should increase with age [19, 20]. Interactions within oligodendrocytes were also shown to decrease with age.

To establish the specific LR interactions that changed significantly in different sender and receiver cell types between age groups, MMCCI’s LR pair network dissimilarity ranking was run on the integrated aged and young samples (**Fig. 4b**). The most dissimilar pair, Vimentin (Vim)-Cd44, showed interactions mostly within and between neurons, astrocytes, and oligodendrocytes in the aged brain, while in the young brain, there were no interactions with astrocytes, but more ependymal cell interactions (**Fig. 4b**). Vim is a known astrocyte marker, expressed in ependymal cells in healthy brains and in reactive astrocytes in aging [21, 22]. Cd44 is an astro-mesenchymal marker, so this interaction was expected to change with age, with Vim release shifting from ependymal to astrocytes during aging [23]. For the second example, the A2m-Lrp1 pair was detected to interact between neurons, oligodendrocytes, astrocytes, and ependymal cells in the aged brain, but had no interactions with astrocytes in the young brain. This LR pair has been shown to potentially associated with Alzheimer’s disease [24, 25].

These differential comparison methods can be applied to any samples, integrated or not, to quantitatively and qualitatively compare different biological groups. Samples can be compared at the whole network level or at the individual LR and cell-type level, providing valuable answers into any important biological questions involving differential analysis.

#### MMCCI queries LR interactions between specific cell-type pairs

To obtain deeper insights into specific cell-type sender-receiver pairs, MMCCI provides functions for querying LR pairs and their proportions involved in specific CCIs and can perform pathway analysis on these pairs. Since the interactions from astrocytes to neurons were significantly increased in the aged brain, this set of interactions was queried to find the LR pairs involved along with their relative interaction strength, quantified by their interaction score proportion in that sender-receiver cell-type pair (**Fig. 4c**). These results show that not only did the overall strength of the interactions between the cell types increase in age, but so did the number of LR pairs as well. We found many different LR pairs interacting in the aged group compared to the young group.

In many of these interactions, predominantly in the aging brain, Lrp1, a multifunctional cell surface receptor, was involved [26]. In the young brain, Midkine (Mdk), a neurotrophic growth factor that is involved in growth and proliferation during embryogenesis, was one of the active ligands, while in aging, the serpin family became the predominant ligands acting on Lrp1 [27]. In the young, another top ligand, Itgav, has been reported to play an important role in neurovascular cell adhesion in brain angiogenesis [28]. This analysis in MMCCI can provide interpretable results to address biological questions involving specific interacting cell types.

#### MMCCI clusters LR pairs with similar cell-type networks

To summarise the large number of interactions, MMCCI provides a LR network clustering algorithm, which in this case was applied on the integrated aged and young samples separately. This analysis identifies and groups LR pairs with similar cell-type networks under the assumption that each cluster of LR pairs are likely to have similar biological roles. Further, Enrichr analysis was run on each cluster to find the pathways involved in the interactions between cell types in a particular cluster (**Fig. 4d**). The selected aged cluster in **Fig. 4d** grouped LR pairs with similar interaction networks predominately in astrocytes, oligodendrocytes, and neurons, and we found pathways related to inflammatory response, complement, not present in the young cluster that are significant components of the astrocyte to neuron and oligodendrocyte interaction in aging [29]. The selected young cluster grouped LR pairs with similar interaction networks in neurons, and discovered neurodevelopmental pathways such as angiogenesis, axon guidance, and neuroactive LR interactions. These results show analyzing a specific cluster of LR pairs can reveal specific and relevant biological pathways.

#### MMCCI identifies LR pairs and cell-type interaction networks involved in specific aging-related pathways

To look deeper into specific biological pathways, MMCCI provides functions to identify and rank the LRs involved in user-specified pathways from Encrichr and create cell-type networks for the given pathways. In this aging dataset, LR pair networks involved in the complement and Wnt signaling pathways were extracted from the aged and young integrated samples respectively and summed, allowing the visualisation of the cell types involved and the relative strengths of the LR pairs involved (**Fig. 4e**). In the aged complement pathway, interactions between astrocytes, neurons, oligodendrocytes were the strongest, with neurons acting overall as more of a receiver. Lrp1, a protein found predominantly on neurons and reactive astrocytes, was the most common receptor in the top complement interactions [30]. This receptor, particularly through interactions with A2m, is known to be involved in Alzheimer’s disease pathogenesis [24]. The young Wnt/*β*-catenin pathway had interactions in oligodendrocytes, ependymal cells, and neurons, which is expected as this pathway is known to have an important role in neuron development and myelin formation [31, 32]. These results demonstrate how MMCCI’s unique and powerful CCI pathway analysis can offer useful insights into the cell types and LR pairs involved in specific biological pathways.

#### MMCCI spatially clusters cells/spots with similar LR interaction scores

MMCCI also offers the functionality to identify spatial communities based on common LR pair interactions by performing LR interaction clustering. Applying this to the aging brain dataset, we can cluster tissue regions similar to standard cell/spot clustering, but using LR interaction scores, which are based on the coexpression of the ligand and receptor in each spot and its neighbouring spots, calculated by stLearn CCI results for each individual sample. This is different to common clustering methods using gene expression. The plots in **Fig. 4f** correspond to the two marked samples in **Fig. 3a, 3b**, where we found the interacting cell-types, and in **Fig. 4f** we can visualise the spatial location where those cell-types are interacting. The stacked bar plots show the proportions of each cell-type in different clusters, revealing in this comparison how there are far more astrocytes in the aged sample and they are involved in many different clusters, meaning they are involved in a wide range of interactions (**Fig. 4f**). These resulting clusters can be used for many other downstream analyses, such as finding marker LR pairs for each cluster. This function allows a new type of clustering to be performed on spatial data, which can provide valuable insights into where certain interactions are occurring spatially.

### MMCCI integration and pathway analysis of multimodal spatial melanoma datasets

We applied MMCCI integration to our multimodal spatial cutaneous melanoma dataset, which comprised ten patients, four of which had Visium and CosMx samples and six with only Xenium samples, in order to collate a set of interactions that are significant across patients and technologies. We then investigated the cell types and LR pairs involved in important melanoma-related biological pathways.

For patients with matched Visium and CosMx data, samples were integrated at patient-level. Notably, individual CosMx and Visium samples demonstrated high intra-sample variability, with cell-cell interactions identified between all cell types. Many of these interactions were identified in only one of the two modalities and were thus likely to be circumstantial rather than biological; averaging these data across regions of interest would lead to distortion of the final dataset. In contrast, patient-level integration through MMCCI enabled exclusion of these ‘false positive’ interactions while retaining the fibroblast, vascular, and immune cell-cell interactions consistently present across all samples (**Fig. 5a**).

**Fig. 5.**
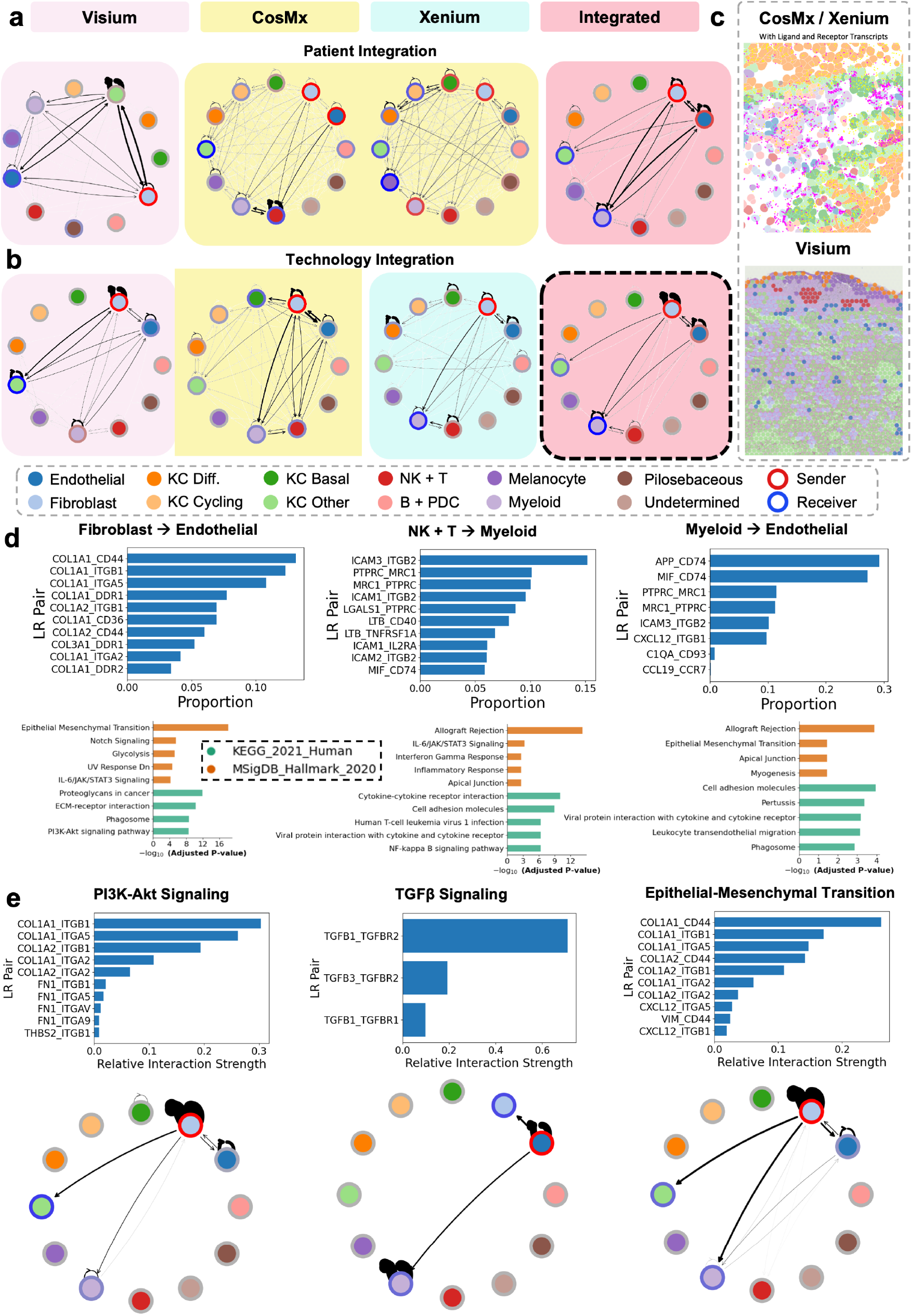
Integration and analysis of multimodal human melanoma CCI results. **a**. CCI integration results for a patient, where one Visium (pink) and two CosMx (yellow) samples of the same patient were integrated together (red). **b**. Within-technology integration results for four Visium (pink), eight CosMx (yellow), and six Xenium (blue) samples, along with between-technology integration of all samples (red). The integrated sample (bordered) is employed for all subsequent downstream analyses. **c**. Two representative melanoma spatial plots, showing the difference between the single-cell resolution image-based technologies, CosMx and Xenium, and the non-single cell spatial sequencing-based technology, Visium. For the CosMx sample, the transcripts for COL1A1 (magenta) and CD44 (yellow) are plotted to show the resolution of these imaging-based technologies. **d**. Three examples of LR pairs and their proportions that are interacting from a selected sender to a receiver cell type pair, followed by pathway analysis of the LR pairs using Enrichr with the KEGG 2021 Human and MSigDB Hallmark 2020 databases. **e**. LR pairs from the integrated sample involved in different cancer-related pathways and the cell-type network of these LRs.

Samples were also integrated at technology-level to generate composite samples/networks for each of the Visium, CosMx, and Xenium modalities; each of these was in turn aggregated using MMCCI into one final integrated sample/network (**Fig. 5b**). This integrated multimodal sample/network was highly similar to the integrated patient-level samples, highlighting the potential of the Xenium samples in validating the other technologies’ findings. This between-technology integrated sample was used for all the further downstream analyses. These integration results highlight the ability of MMCCI’s multimodal integration pipeline to pull a significant set of interactions out of an inconsistent or noisy set of CCI results.

#### MMCCI provides LR pairs and biological pathways involved in specific cell types

To understand the biological processes involved in the interactions occurring between particular cell types of interest, we analysed the Enrichr pathways of the LR pairs interacting across three different sender-receiver cell-type pairs (**Fig. 5d**). Enrichment analysis identified a high level of interactions between tumour-associated fibroblasts and endothelial cells, involving the COL1A1 and COL1A2 ligands. These interactions, characterized by the binding of Type I collagens to endothelial CD44 and ITGB1 receptors, are key mediators of ECM remodelling, which in turn facilitates melanoma cell invasion and epithelial-mesenchymal transition (EMT) [33, 34]. Correspondingly, pathway analysis demonstrated upregulation of gene pathways representing focal adhesion, ECM-receptor interaction, and epithelial-mesenchymal transition in the fully integrated sample/network (**Fig. 5d**).

Interactions from natural killer and T cells to myeloid cells included the leukocyte adhesion molecules ICAM3 and ITGB2, the NF-*κ*B activators TNF1A and LTB, and the MIF-CD74 LR pair [35]. The pathway analysis consequently indicated upregulation of cell adhesion molecule signalling and the pro-inflammatory JAK-STAT and NF-*κ*B pathways (**Fig. 5d**), highlighting the capacity of MMCCI to capture the immune response to melanoma at spatial resolution and with respect to specific cell types. MIF-CD74 remained the primary ligand-receptor pair for myeloid-endothelial cell interactions, and this interaction was previously demonstrated to promote melanoma cell survival through IL-8 and ERK-mediated signalling [36]. Pathway analysis demonstrated upregulation of leukocyte migration and adhesion pathways, as well as EMT in the context of enriched PI3K-Akt signaling (**Fig. 5d**). These results show how analyzing specific cell-type interactions can provide insights and important hypotheses about the cell-cell interactions in specific tumours that may be obscured when considering only the bulked or global interactions in a sample.

#### MMCCI identifies LR pairs and cell-type networks involved in specific melanoma pathways

To further interrogate the LR pairs and cell-types involved in specific cancer-related pathways, we performed a subset analysis on integrated CCI results corresponding to three selected pathways. Enrichr pathway analysis was applied to specify the LR pairs involved in each included pathway. The PI3K-Akt signaling pathway was characterized by interactions between Type I collagens and integrin receptors (**Fig. 5e**), indicating a CAF-driven milieu that predisposes to ECM remodelling and consequent EMT [37, 38]. This was complemented by the TGF-*β* signaling pathway, which involved enrichment of the TGFB1-TGFBR2 LR pair across multiple cell types: both homotypic endothelial and myeloid cell interactions as well as sender-receiver interactions from endothelial cells to fibroblasts and myeloid cells (**Fig. 5e**). These findings align with the established roles of TGF-*β* signaling in promoting EMT and inducing differentiation of tumour-associated macrophages. [39, 40].

For LR pairs implicated in EMT, the COL1A1 and COL1A2 ligands and the CD44, ITGA1, ITGA5, and ITGB1 receptors were most frequently represented, as were the mesenchymal proteins VIM and FN1 (**Fig. 5e**). The majority of interactions were between fibroblasts to keratinocytes, myeloid cells, and endothelial cells, which corroborates the established contribution of cancer-associated fibroblasts (CAFs) to pro-EMT signaling in the tumour microenvironment (**Fig. 5e**) [41, 42]. This type of analysis provided by MMCCI can be applied with any pathway related to the user’s dataset and is able to expand on current pathway analysis pipelines by generating a cell type network and ranking the LRs by interaction proportion for a given pathway.

### Investigating integration performance across CCI methods using simulated data

To benchmark MMCCI’s performance across multiple CCI tools, both spatial and non-spatial, we integrated three simulated breast cancer spatial samples that were run through four different CCI methods - stLearn, CellChat, NATMI and Squidpy (**Sup. Fig. 2**). For each CCI tool, the integrated result was able to compile a clearer picture of the interactions as a whole across the samples, where consistent interactions are made stronger and inconsistent interactions are made weaker in the integrated results.

Each sample had cell types with neighbors common across all the samples as well as neighbors unique to that sample. This allowed us to confirm whether integration could recover the known ground truth from the simulated data, which were the cell types adjacent to each other in the majority of the samples (for example, B and T cells). This approach also improved detection and exclusion of false positive interactions, such as cell types that were proximate in only one of the samples and were distal/exclusive in remaining samples (e.g., endothelial and stromal cells). In both simulated and biological data, we observed that MMCCI successfully removed the false positive interactions while retaining the true interactions across several spatial and non-spatial CCI methods. We also found that using stLearn CCI on multiple samples and integrating with MMCCI resulted in the highest performance with detecting expected biological interactions while filtering false interactions [3], an observation that can be attributed to the underlying algorithm that stLearn uses, which utilizes spatial information in the analysis to find more confident interactions.

## Discussion

MMCCI was developed as a fast, comprehensive, and open-source Python package for integrating cell-cell interactions from multimodal transcriptomics datasets, along with a downstream analysis toolset for exploring the biological interactions in single samples as well as MMCCI-integrated results. MMCCI is the first platform that enables integration of CCI networks from multiple samples and modalities, strengthening the concordance of the inferred interactions. MMCCI integrates, analyzes and visualizes outputs from multiple CCI tools, including stLearn, CellPhoneDB, Squidpy, CellChat, and NATMI. Recently, CellCommuNet was developed to integrate scRNA-seq samples before running CCI analysis, but unlike MMCCI, it does not integrate the CCI results themselves and is also limited to non-spatial scRNA-seq samples [43]. LIANA is a different tool that runs multiple CCI methods on a single sample and calculates a consensus rank for interactions [44]. However, this is also different to MMCCI in that it only processes a single scRNA-seq sample rather than integrating different samples together.

Validating MMCCI’s integration method on spatial single-cell whole-transcriptome CosMx human pancreas data by comparing the MMCCI integrated CCI results of each FOV to the CCI results of the whole sample, we showed that MMCCI’s integration is significantly more accurate to the ground truth than averaging the CCI results across the FOVs (**Fig. 2**). Not only were CCIs common across the FOVs integrated accurately, but also rare CCIs with high variation across FOVs (**Sup. Fig. 1**). Benchmarking the integration method on simulated spatial breast cancer samples on four different CCI methods showed that MMCCI is able to successfully integrate CCI results from both spatial and non-spatial methods and preserve important cell-type interactions across the three samples (**Sup. Fig. 2b**). Applying MMCCI’s integration and downstream analyses pipeline on stLearn CCI results from multimodal brain and melanoma data, we show the effectiveness of using MMCCI to statistically integrate CCIs both within and between technologies. MMCCI accurately identifies the frequency and distribution of canonical mediators and cell-cell interactions involved in specific biological pathways by integrating CCI results across multiple samples and modalities. Data from each individual sample harbours some insight into the true biological cell-cell communications, but analyzing a single sample is prone to producing false discoveries or missing some important information. This advantage of MMCCI integration is clearly seen in the melanoma single-patient integration, shown in **Fig. 5a**. Individually, the CosMx samples report putative interactions between all cell-types, many of which are likely falsely detected or not biologically relevant. While the Xenium samples are more targeted, they omit a number of expected interactions, such as those between natural killer (NK) cells, T cells, and melanocytes. Integration resolves this discrepancy to generate a clear, consistent, and biologically pertinent profile of the interactions underlying thin primary melanoma. Through the integration of multiple CCI results from different samples using MMCCI, we can gain a deeper understanding with higher confidence about the intercellular interactions occurring in normal physiology and disease.

MMCCI was able to provide relevant biological insights into the multimodal aging mouse brain and melanoma datasets that may have been overlooked if only one sample or transcriptional modality was analyzed. A clear example of this is that biologically important interactions with vascular cells are missing in many of the Visium brain samples, but after integration with STOmics samples, these are preserved (**Fig. 3**). Many important interactions in the aging brain were observed in the integrated results. Mainly, astrocytes were shown to become more active, not only increasing their overall interaction strength but also the LR pairs involved in the interactions, which is consistent with the current understanding of the roles of astrocytes becoming more reactive in aging, contributing to neuroinflammation and the complement pathways [45, 19, 20, 21, 22, 23, 46, 47, 48]. The thinning of the ependymal layer expected in aging was also shown by the observed decrease in ependymal interactions in aging [17, 18]. As well, many neuron development pathways were observed in the young samples which were not present in the aged. Overall, MMCCI allows for deep insight to the molecular mechanisms involved in aging in the central nervous system.

In primary cutaneous melanoma, MMCCI was able to synthesize data from complementary ST technologies to delineate the contributions of Akt-mediated extracellular remodelling and TGF-*β* signalling in promoting epithelial-mesenchymal transition within the stromal milieu [49]. Single-sample analyses captured interactions across all cell types, restricting interpretation of the findings; however, MMCCI filtered through these to obtain consistent CCIs across the melanoma dataset. Notably, MMCCI highlighted the broad activity of fibroblast-associated Type I collagen ligands across multiple receptors and gene pathways, including targeting of both endothelial and myeloid cells [50, 51]. Another example of a pertinent CCI observed in the integrated sample involves the macrophage migration inhibitory factor (MIF)-CD74 LR pair. MIF-CD74 interaction promotes melanoma cell survival through recruitment and activation of the PI3K/Akt signalling pathway, and it is a promising candidate for targeted therapy [52, 53, 54]. Overall, MMCCI enabled precise exploration of the complex network of interactions within the tumour microenvironment, particularly with respect to the role of fibroblasts in ECM remodelling, EMT, and PI3K-Akt signaling. This highlights the utility of the MMCCI pipeline in meta-analysis of multi-sample and multi-modality data to infer biologically significant interactions with a higher degree of confidence than from any one sample or modality alone.

While MMCCI offers a range of new and important, there are limitations that need consideration during implementation. Imaging-based technologies often detect fewer interactions that would be expected biologically as a result of having a lower number of proteins/genes in the panel. Because of this, a future direction of CCI result integration could explore the option to impute missing genes/proteins. However, this is not always a significant limitation as the gene panels are often selected based on biological relevance to the type of sample, so the integrated results still provide valuable insight into the relevant biological pathways. Alternatively, users can also focus on data integration for just the genes/proteins that are shared across platforms. Users can also select parameters to control for the integration weights between modalities that take into account the omission of genes/proteins in the panels. Meanwhile, sequencing-based technologies can measure more genes, but are often not yet able to provide single-cell resolution. The low resolution means that while many LR pair interactions are able to be inferred, the cell-types involved are less specific and lack the precision of imaging-based technologies.Through the integration of CCI results from multiple samples, MMCCI is able to use the advantages of both imaging- and sequencing-based spatial transcriptomics to construct a clearer picture of the biological interactions across multiple samples and technologies.

Overall, the MMCCI package provides novel integration method for statistical meta-analysis of multi-sample or multimodal CCI results. MMCCI’s flexible integration pipeline allows for samples to be integrated more strictly to filter to only highly confident interactions across the samples, or for the integration to include rarer interactions that are significant in only a subset of the samples, which makes the pipeline robust to cancer samples where there is tumour heterogeneity between samples and important interactions are not removed due to being rare. MMCCI is the first tool that can integrate both spatial and non-spatial CCI results from different transcriptomic modalities and CCI methods together, and will prove relevant to future analyses in the rapidly evolving field of spatial transcriptomics.

## Conclusion

We have developed a new algorithm, MMCCI, which is the first tool for statistical integration and meta-analysis of multimodal spatial transcriptomics and scRNA-seq CCI results from different tools. We have benchmarked MMCCI integration pipeline on spatial single-cell whole-transcriptome human pancreas data and verified that its accuracy outperforms simply averaging CCI results across samples. MMCCI integration and downstream analysis was applied to multimodal brain and melanoma spatial transcriptomics datasets to uncover the cell-types and LR pairs involved in complex aging and cancer related pathways. We provide the MMCCI software as an open-source package and have included detailed usage instructions, aiming to facilitate robust CCI analyses in the broader community.

## Methods

### 1. Upstream data processing

#### 1.1. Deconvolution

For the aging brain samples, we generated Visium and STOmics data and annotated cell types using RCTD with the Allen Brain Atlas as a single cell reference for deconvolution [55, 56]. For the melanoma samples, we generated Visium, CosMx and Xenium data, followed by automated cell-type annotation by deconvolution and label transferring using an in-house single cell skin cancer reference. Specifically, for Visium data, we used RCTD for finding cell-type compositions of spots which contain 1 - 10 cells, and for the single cell resolution CosMx and Xenium data, we used Seurat V4 Label Transfer for direct single cell to single cell annotation [57].

#### 1.2. stLearn cell-cell interaction analysis

Cell-cell interactions for the pancreas, brain and melanoma samples were computed using stLearn, a CCI inference method that takes spatial information into account [3]. We first filtered out genes that were not present in more than three spots and performed counts per cell normalization. stLearn performs ligand-receptor analysis, in which each spot or cell receives an interaction score from its immediate neighboring spots for each LR pair. The LR pair database used in stLearn was from connectomeDB2020 [7], but other databases can also be applied. This score is used for MMCCI’s LR interaction clustering. Importantly, stLearn also predicts significant cell-cell interactions, generating a network of communicating cell-types represented as a matrix per LR pair where the weights are the number of significantly interacting spots or cells, along with a p-value for each weight. This data is used by MMCCI to perform integration and is the format used by most of the integration, analysis and visualization functions. CCI results obtained from other methods are converted to this format by MMCCI for compatibility with the package.

### 2. Integration method

Sample integration (**Sup. Fig. 3**) is split into two sections, within-technology and between technology integration, which can both be run separately.

#### 2.1. Within-technology integration

This step performs CCI integration of samples within a technology by accounting for differing sample sizes, reflected by the number of spots/cells in each sample (1.1 of **Sup. Fig. 3**). Each sample’s LR pair interaction matrices are scaled based on the number of spots/cells in that sample (sample-size scaling). Each weight in each LR matrix is multiplied by this scaling factor. This is required to control for samples with a disproportionate number of spots, ensuring that each sample has an equal weighting within each technology as a sample with more spots will generally have more interactions.

An integrated sample for a given technology is then created by running a LR-level integration function (**Equ. 1 - 2**) with each sample arising from that technology (1.2 of **Sup. Fig. 3**). The CCI integration works by first creating a list of LR pairs to be included in the integrated sample by either taking the common LR pairs if there are two samples to integrate, or by taking the LRs present in at least half the samples if there are more than two samples to integrate (**Equ. 1**). The option is given in the package to select other methods for selecting which LR pairs to integrate including using LR pairs present in all samples, the majority of samples, or at least one sample. Each LR pair in sample *k* has a cell-type adjacency matrix which has an interaction score for that LR pair for cell-type *i* to cell-type *j*, represented as 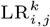 and referred to in this paper as an interaction. Then for each LR pair in that list, the integrated network for that pair is calculated by combining the networks of each sample, creating a matrix where each interaction is the geometric mean of that interaction in all the samples where the score is not 0 (**Equ. 2**). If more than half of the samples have an score of 0 for a given interaction, it will have an integrated score of 0 for that interaction.

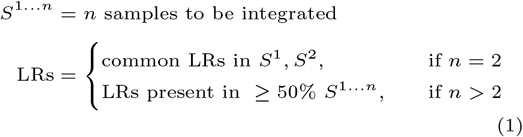

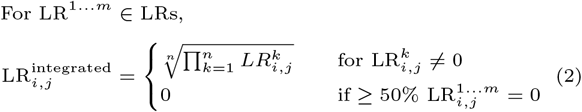

Non-significant interaction scores in each LR matrix are then set to zero. The p-value of an interaction for a sample is provided from the CCI analysis performed on the sample, and the integrated p-value across all samples for an interaction of an LR pair between two cell-types is calculated as outlined in section 2.3 of the Methods.

#### 2.2. Between-technology integration

The integrated CCI values for each sample of each technology are then normalized by a scaling factor, calculated by averaging the arithmetic mean of the values across all the LR pairs per technology (2.1 of **Sup. Fig. 3**). This ensures that a high number of interactions detected through one specific technology are not over-represented in the integrated sample.

A final integrated sample is then created by running the LR-level integration function as in the within-technology integration (2.2 of **Sup. Fig. 3**). After this, the overall network of interactions, an overview of the cell-type interactions in the sample, is calculated by scaling the values in each LR matrix so that they sum to one to ensure equal weighting of all LR pairs, and then computing the mean of all the matrices. Creating an overall network can also be done in other points of the integration to easily visualize the overall interaction network of an integrated or non-integrated sample.

Again, the p-values across all samples used in the integration can be combined as outlined in section 2.3 of the methods. These p-values can be used to filter insignificant interactions in the integrated sample.

#### 2.3. Meta-analysis: a statistical framework for integration

Meta-analysis is a statistical technique to combine the results from multiple different studies addressing a similar question. Most of the CCI algorithms that MMCCI is able to integrate provide p-values for every interaction score, which are integrated to calculate a final p-value for each interaction in the integrated results. We used Stouffer’s method to integrate p-values from CCI results of different samples, which converts p-values to z-scores before combining them together [58, 59]. The z-scores are calculated using the inverse cumulative distribution function of p-values (**Equ. 3**), integrated (**Equ. 4**), and then converted back into a p-value using the cumulative distribution function (**Equ. 5**). This meta-analysis improves the robustness and accuracy of MMCCI’s integration method.

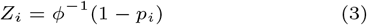

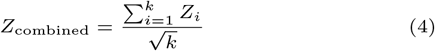

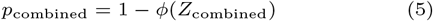

### 3. Downstream analyses

#### 3.1. Dissimilarity scoring

The dissimilarity score shown in (**Sup. Fig. 3** and **Equ. 6**), has been based off of a similar metric introduced in COMUNET [10]. To compute a dissimilarity score between two matrices, we first ensure that they have the same interacting cell types by imputing an interaction score of zero to the missing cells in the matrix with fewer cell types. For any two cell-type adjacency matrices, *M*_1_ and *M*_2_, the continuous dissimilarity score, *S*, is calculated by taking the absolute difference between the weights of each sender-receiver cell-type pair of the two matrices divided by the sum of the weights. The binary dissimilarity score, *F*, is calculated as 1 if the two cells of the matrices have a different number of significantly interacting spots, or otherwise 0. Finally, these two dissimilarity scores are added for each LR-pair using a blending factor, *λ*, and this score is divided by the square of the number of cell-types, giving the dissimilarity score, *D. D* is an approximate representation of the proportion of the network that is different, and for most cases this difference is around 0 − 0.2 due to many edges having a weight of 0 in both networks as it is rare to find LR pairs that interact between all cell-types in a sample.

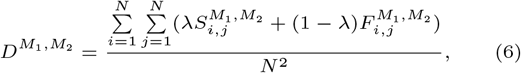

For *S* (**Equ. 7**), the denominator adds the 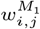 and 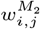 and these are subtracted in the numerator, due to which S is lower for interactions with same numerator, where both values are non-zero compared to one of them being zero. So, *F* (**Equ. 8**) helps in increasing *D* in cases where there are many non-zero interactions, as *S* will be lower for two non-zero values with the same difference as two values where one is zero. This can lead to *D* being lower than expected in comparison to two networks each with only a few interactions. By default, we set *λ* = 0.5 to equally weigh the two scoring metrics. The effects of changing *λ* across different LR pairs is shown in **Sup. Fig. 2c**. For networks with many edges that change slightly, the dissimilarity is higher when *F* is weighted higher, whereas in networks where there are some edges in one and none in the other, there is no difference between *F* and *S*.

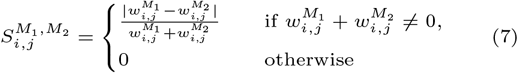

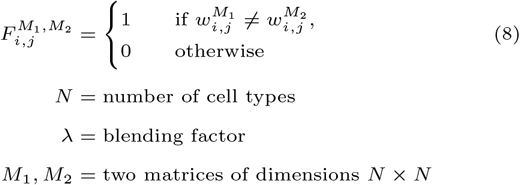

#### 3.2. Differential CCI permutation testing

CCI results provide a network of the different interacting cell-types. The edge represents the number of cells/spots involved in the interaction. Here, the permutation test is used to find significantly different interactions between two networks. The overall integrated interactions for samples across two groups are first normalized separately so that the interaction count matrix sums to 1 and then the difference between the two matrices is calculated. Next, permutation testing is performed by repeatedly shuffling the values along both rows and columns, and then observing how often the differences in shuffled matrices are bigger than the true differences, returning a p-value to identify which specific interactions between two cell-types are statistically higher than the background. This can also be done instead to calculate the differences between any two networks, such as a comparing a single LR pair between two different groups or comparing between two different LR pairs in a sample.

#### 3.3. LR network clustering

For the LR network clustering represented in **Fig. 1b**, we calculated a pairwise dissimilarity score matrix for each LR pair (**Equ. 6 - 8**). Using this matrix, we calculated a distance matrix using “pdist” and “squareform” from the SciPy package [16]. The distance matrix was normalized using min-max normalization. We then computed the PCA of the dissimilarity matrix and used the first two components along with the distance matrix for clustering of the LR networks. This function allows the user to perform either hierarchical or k-means clustering, and finds the optimal number of clusters unless stated otherwise.

#### 3.4. LR interaction clustering

This method is only currently available for individual ST samples processed through stLearn, but can be applied to CCI results for any method that gives an LR interaction matrix for each spot. LR interactions were scored from stLearn, which give a score for each LR pair for each spot. The calculation was based on the interactions of LRs within a spatial distance up to 250*µ*m. Shown in **Equ. 9**, stLearn calculates a location-specific LR score for each spot, *S* with spatial neighbours, *N* [3].

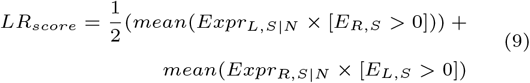

LR interaction clustering helps to identify groups of cells that are interacting through similar LR pairs. These scores are pre-processed and clustered using the Scanpy spatial transcriptomics analysis pipeline for Leiden clustering [60]. Each cluster denotes a spatial interaction module involving interaction of multiple cell-types, formed based on LRs having similar interaction scores. The stacked barplot indicates the proportion of the interacting cell-types in each cluster. An example is shown in **Fig. 4f**.

#### 3.5. Cell-cell LR pair querying

LR pairs can be queried given a sender and receiver cell type and then ranked based on the proportion of the total number of interactions for the given sender and receiver. This is represented in **Fig. 1f**.

#### 3.6. Pathway analysis

Pathway analysis is performed using GSEApy, a Python package that can perform gene set enrichment analysis using Enrichr [61]. LRs used for analysis are split into the ligand and receptor and are added to a gene list to be run through GSEApy’s Enrichr pipeline, which can be used with any specified Enrichr databases. In this pipeline, both mouse and human genes can be inputted and will be automatically accounted for. Individual pathways can be further analyzed by subsetting a CCI result to only LR pairs that have at least one or both genes involved in the pathway, and then a cell-type network can be plotted as well as the LRs involved and their relative proportion.

### 4. Visualizations

#### 4.1. Network visualizations

Network graph plotting is performed using NetworkX, examples of which are in **Fig. 3** [62]. Nodes have an outer ring color scale based on whether that cell type is sending (red) or receiving (blue) more interactions. When plotting a network difference plot, p-values from the permutation testing can also be given and edges with significant p values are plotted as purple if they are higher in the first group or green if lower, along with the p-value shown on the edge. As an alternative visualization, network chord plots are included and were adapted from the stLearn package [3].

### 5. CCI output conversions

For broad applications, MMCCI has wrapper functions included in the package that allow conversion of multiple different CCI pipelines. Currently, stLearn, CellPhoneDB, Squidpy, CellChat, and NATMI outputs can be converted to be compatible with MMCCI’s pipeline and samples from different CCI methods can be integrated together. The MMCCI pipeline is outlined in **Fig. 1** and **Sup. Fig. 3**.

### 6. In-house spatial transcriptomics data generation

The brain and melanoma datasets were generated in-house, while the CosMx Human Pancreas FFPE Dataset was a publicly available dataset from NanoString and the scRNA-seq breast cancer was sourced from a study by Karaayaz et al. that was publicly available [63].

#### 6.1. Brain samples

Young (3 months) and aged (20–24 months) C57BL/6 mice (all female and housed in the same animal facility) were sacrificed for organ harvest between 8am and 9am to avoid molecular changes related to differences in circadian rhythm. Only healthy animals were processed; mice bearing tumors or displaying any internal alterations (e.g., enlarged spleen, inflamed liver etc.) were excluded.

For Visium spatial gene expression, the tissue sections were placed on the pre-equilibrated Visium Spatial Tissue Optimisation Slide (10X Genomics, cat no.3000394) and Visium Spatial Gene Expression Slide (10X Genomics, cat no.2000233). The Visium library was constructed according to the Visium Spatial Gene Expression User Guide (CG000239 Rev B, 10X Genomics).

For STOmics, the tissue sections were placed on Stereo-seq chips (BGI STOmics, cat no. 211SP118). The tissue permeabilization was optimized according to STOmics Stereo-seq Permeabilization User Manual (Version A0). The STOmics library preparation was carried out as described in STOmics Stereo-seq Transcriptomics User Manual (Version A0).

### 6.2 Melanoma samples

Six formalin-fixed, paraffin-embedded (FFPE) primary cutaneous melanoma samples were obtained from a matched case-case series of human patients with thin melanoma [64] for downstream Xenium spatial transcriptomics. Tissue preparation for Xenium in situ gene expression was carried out as described in the Tissue Preparation Guide (CG000578 Rev C, 10X Genomics). In short, six 5 *µ*m FFPE sections were multiplexed on one Xenium Slide (10X Genomics, cat no.3000941) to maximize the usage of the capture area while minimizing batch effect. The detailed onboard image processing, decoding and cell segmentation protocols have been described in Janesick. et al., 2023 [13].

An additional eight primary melanoma samples from four patients were selected for single-cell CosMx RNA sequencing, which were processed by NanoString; of these, four had adjacent sections available for concomitant Visium Spatial Gene Expression.

### 7. Benchmarking

#### 7.1. Benchmarking on pancreas data

The CosMx Human Pancreas FFPE Dataset from NanoString was used to benchmark MMCCI’s integration method as an example with single-cell whole transcriptome spatial data. This dataset consisted of 18 field of views (FOVs), but could also be combined together into one sample. Because of this, both the whole sample and each FOV individually were processed through the standard stLearn CCI pipeline, and MMCCI was used to integrated the CCI results of each FOV. The integrated results from MMCCI were then compared to the CCI results of the whole sample, which was the ground truth, to quantify the accuracy of MMCCI’s integration method. This pipeline is shown in **Fig. 2a**.

The similarity score used as the metric in **Fig. 2c** was calculated for each LR pair by comparing the CCI matrices of both samples to the whole. This was calculated by taking the proportion of the number of cells in the matrices that there either was or wasn’t an interaction commonly in both matrices.

#### 7.2. Benchmarking on simulated breast cancer data

The simulated spatial breast cancer data was generated using a scRNA-seq breast cancer dataset. The gene expression distribution in each of the cell types was estimated by fitting a negative binomial distribution, and this was used to generate 10,000 simulated cells of each cell type [65] (**Sup. Fig. 2a**). The spatial data was generated by pooling cells of specific types together to simulate Visium data and in a way such that they followed a set of rules about which cells should and should not neighbour (**Sup. Fig. 2b**). These three simulated spatial samples, each with a set of cell types that did and did not neighbor, were then run through stLearn, CellChat, NATMI, and Squidpy CCI analysis and then integrated using MMCCI (**Sup. Fig 2b**). All CCI tools used connectomeDB2020 as the LR database [7]. The individual and MMCCI integrated cell-type interaction networks were generated for the HLA-B:KIR2DL3 LR pair to show how the integration method works at the LR pair level (**Fig. 1c**).

## Declarations

### Ethics approval and consent to participate

The mouse brain tissue experiments were reviewed and approved by The University of Queensland’s Animal Ethics committee and conducted in compliance with the specified ethics regulations. The human melanoma samples were approved for research use under ethics approval numbers 2018000165 and 2017000318 by the University of Queensland’s Human Research Ethics Committees and 11QPAH477 by the Metro South Human Research Ethics Committee.

### Consent for publication

Not applicable.

### Availability of data and materials

MMCCI is available as an open-source Python package. Installation instructions, tutorials, CCI outputs used to produce all manuscript figures, and source code are publicly available via: https://github.com/BiomedicalMachineLearning/MultimodalCCI. The CosMx Human Pancreas FFPE Dataset is publically available from NanoString at https://nanostring.com/products/cosmx-spatial-molecular-imager/ffpe-dataset/cosmx-smi-human-pancreas-ffpe-dataset/. The breast cancer scRNA-seq data is publically available from Karaayvas et al. at https://www.ncbi.nlm.nih.gov/geo/query/acc.cgi?acc=GSE118390 [63].

### Competing interests

The authors declare that they have no competing interests.

### Funding

This work was supported by the Australian Research Council [grant number DE190100116 to Q.H.N.], National Health & Medical Research Council [grant number 2001514 to Q.H.N.], NHMRC Investigator Grant [grant number GNT2008928 to Q.H.N.], and the UQ Genome Innovation Hub, and a MGI DNBSEQ MPS Research Grant awarded to C.M.N..

### Authors’ contributions

Q.H.N., L.H., O.M., conceived experiments and developed the algorithms. L.H. and O.M., wrote the software. A.X. conducted experiments. L.H., O.M., and Q.H.N. analyzed data. K.K., C.M.N., and S.X.T. provided samples and helped with data generation. Q.N., K.K., and C.M.N. provided funding for the work. L.H., O.M., S.X.T., and Q.H.N. wrote the manuscript. All authors have reviewed and approved the manuscript.

## Acknowledgements

We thank our collaborators at the Genome Innovation Hub, Nanostring, BGI and the Frazer Institute for assistance with data generation. We thank all members in Nguyen’s Genomics and Machine Learning Lab. We thank The University of Queensland for providing a scholarship to L.H. and O.M.. We wish to acknowledge The University of Queensland’s Research Computing Centre (RCC) for its support in this research.

---

**Sup. Fig. 1.**
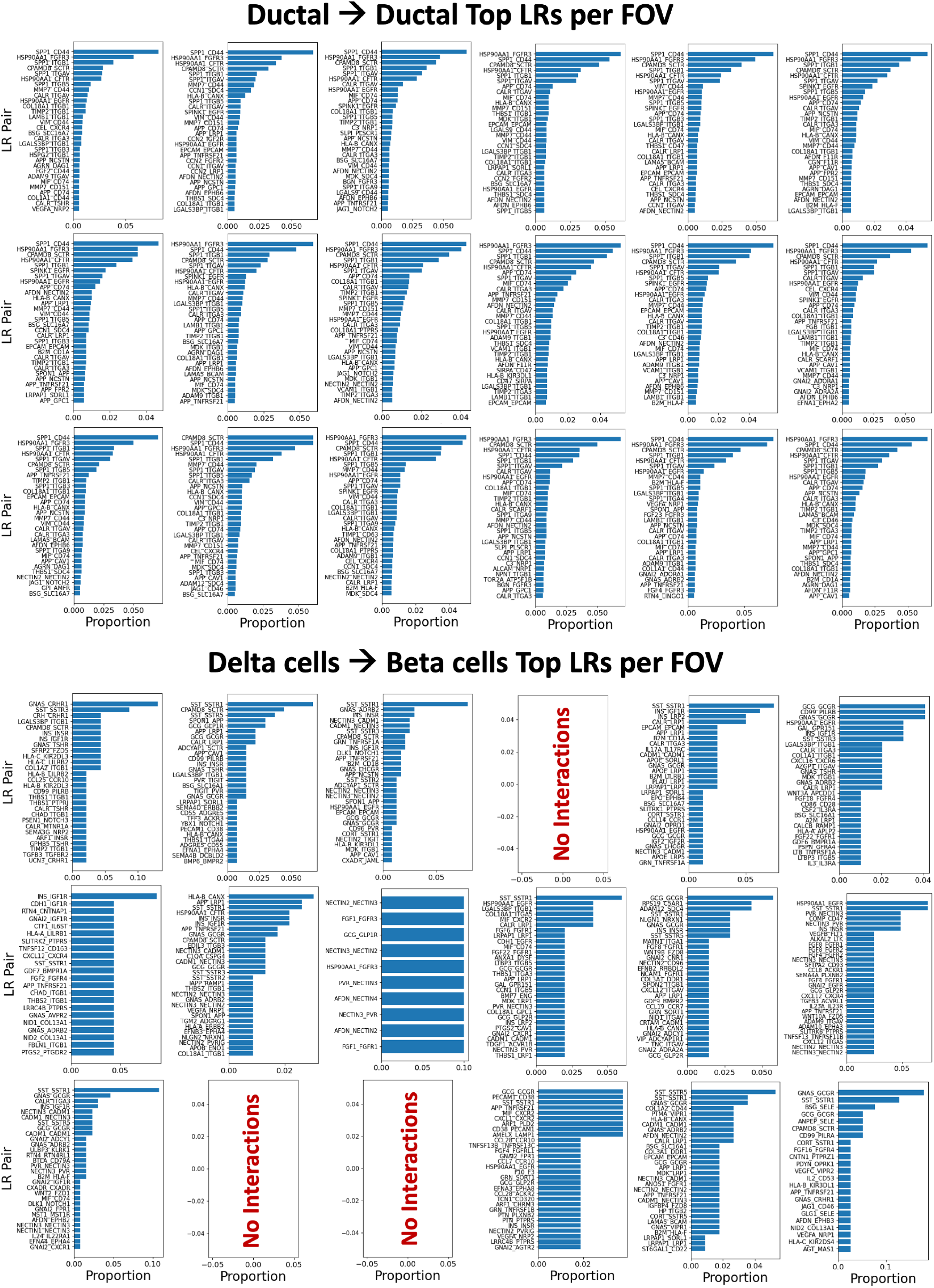
Top LR pairs and their proportions interacting in each of the individual human pancreas FOVs between ductal cells (low variation between FOVs), and from delta to beta cells (high variation between FOVs). The results of all FOVs integrated are shown in **Fig. 3d**.

**Sup. Fig. 2.**
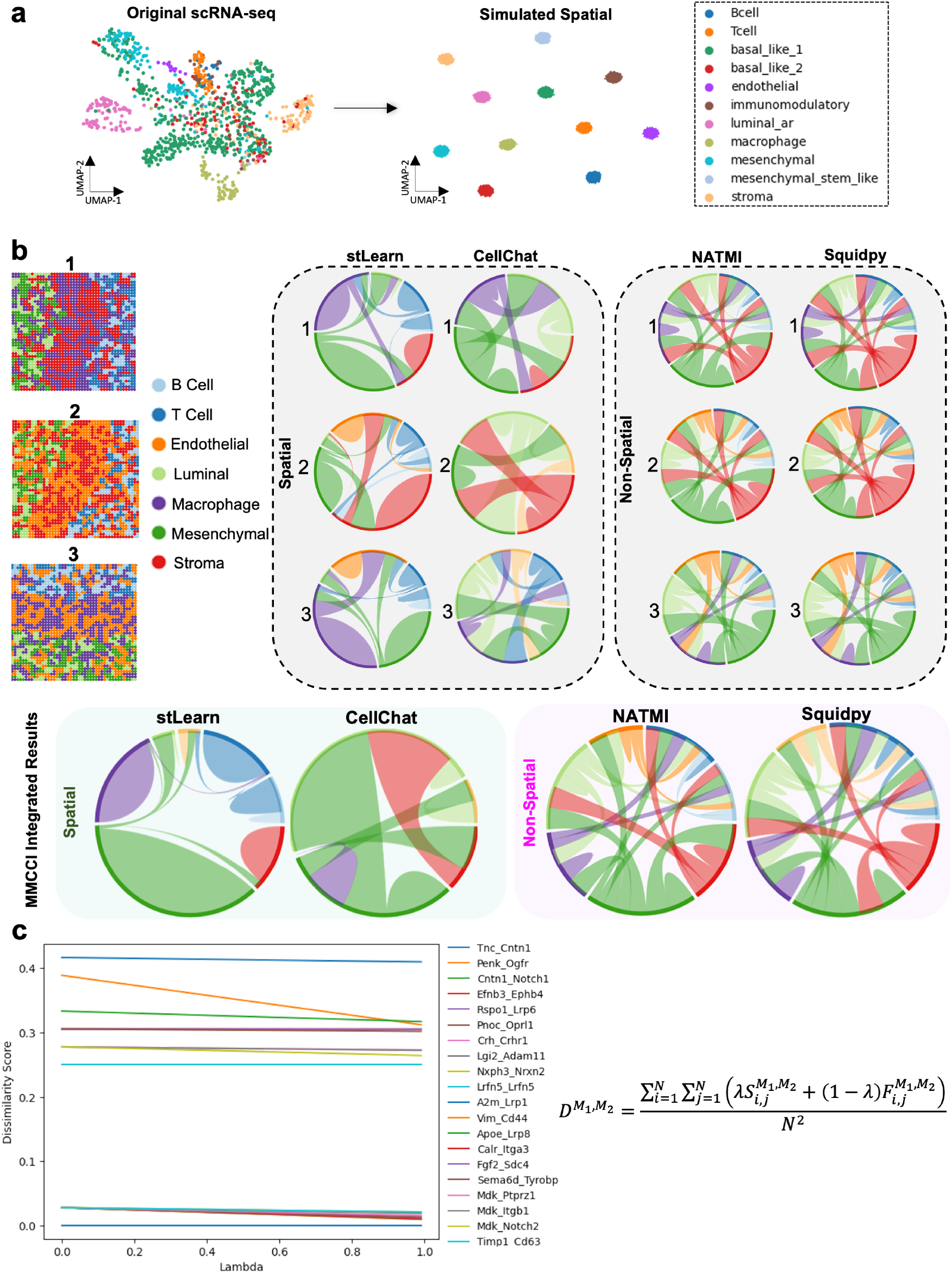
Simulated benchmarking data and dissimilarity score visualization. **a**. UMAP plots showing cell types in the original scRNA-seq and the simulated spatial breast cancer data. **b**. Spatial plots of simulated spatial breast cancer samples (left). Benchmarking results of running within-technology integration on simulated spatial breast cancer samples (1, 2, 3) that have been run through different spatial (stLearn and CellChat) and non-spatial (NATMI and Squidpy) CCI methods. All simulated samples have B and T cells spatially separated from luminal and mesenchymal cells (left). Each sample has a unique combination of two out of stromal cells, macrophages, and endothelial cells. Sample one excludes endothelial cells, sample two macrophages, and three stromal cells. Chord plots show the integrated (left) and unintegrated (right) results for each technology and are used to show the cell types that are interacting. **c**. Line plot showing how changing the value of lambda in the dissimilarity score function impacts the value of the dissimilarity score across different LR pair examples (formula is shown).

**Sup. Fig. 3.**
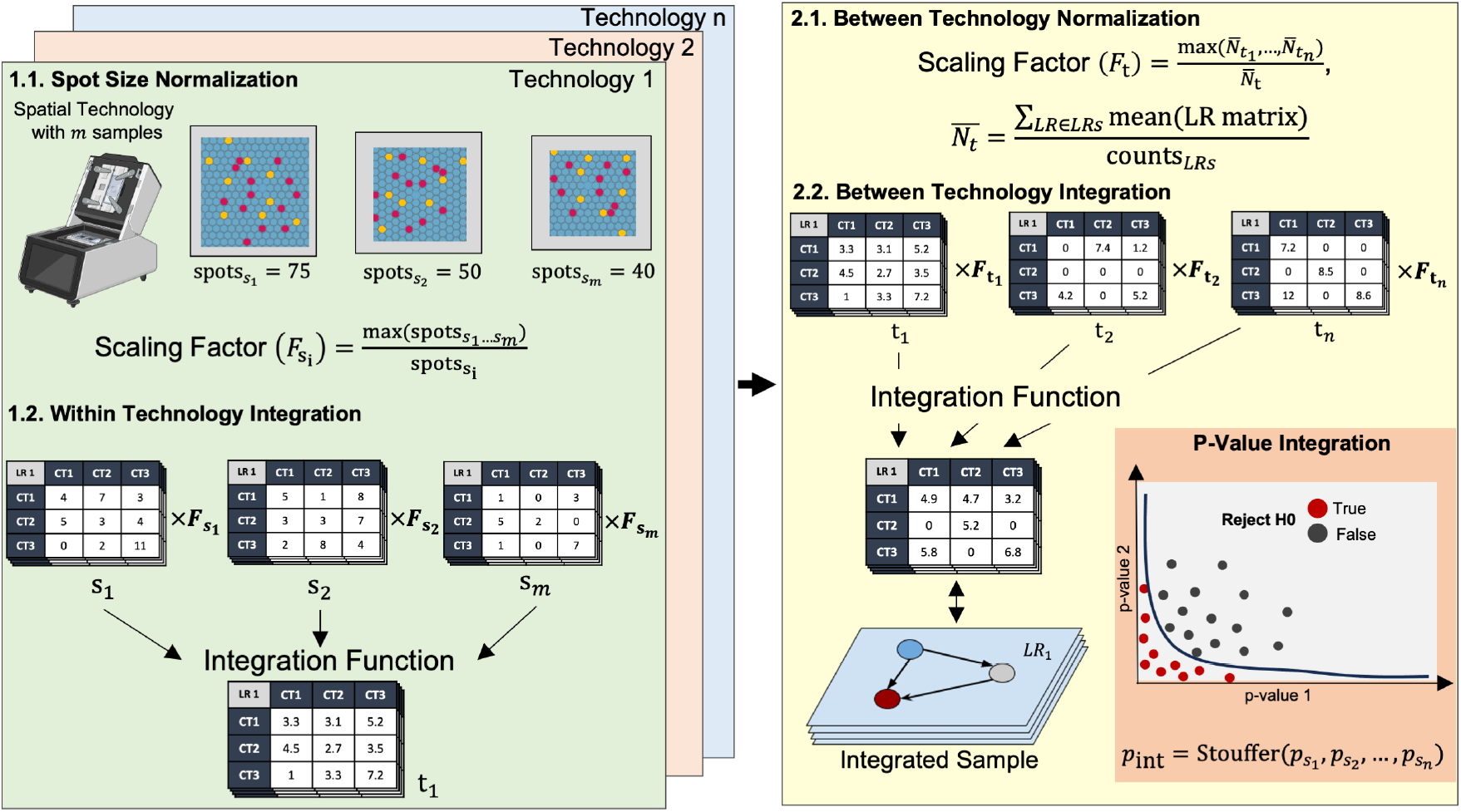
Overview of the integration pathway showing pipeline for integration within (left) and between (right) technologies. 1.1 shows the within-technology normalisation based on number of cells/spots (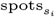) to get a scale factor (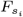). 1.2 shows the integration within a technology. Stacked tables indicate a sample, which is made up of a set of LR matrices. 2.1. shows the normalization between technologies, by calculating a technology scale factor (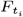). 2.2 shows the integration between technologies into a final integrated sample. LR cell-type interaction p-values from each individual sample (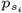) are integrated using Stouffer’s method. Created with BioRender.com.

## Notes

https://github.com/BiomedicalMachineLearning/MultimodalCCI

### Competing Interest Statement

The authors have declared no competing interest.

### Summary of Updates

Many figures revised and new method validation methods added.

## References

1. Bilal N. Sheikh, Olga Bondareva, Sukanya Guhathakurta, Tsz Hong Tsang, Katarzyna Sikora, Nadim Aizarani Sagar, Herbert Holz, Dominic Grün, Lutz Hein, and Asifa Akhtar. Systematic identification of cell-cell communication networks in the developing brain. iScience, 21:273–287, 2019.

2. Xinyi Wang, Axel A. Almet, and Qing Nie. The promising application of cell-cell interaction analysis in cancer from single-cell and spatial transcriptomics. Semin Cancer Biol, 95:42–51, 2023.

3. Duy Pham, Xiao Tan, Brad Balderson, Jun Xu, Laura F. Grice, Sohye Yoon, Emily F. Willis, Minh Tran, Pui Yeng Lam, Arti Raghubar, Priyakshi Kalita-de Croft, Sunil Lakhani, Jana Vukovic, Marc J. Ruitenberg, and Quan H. Nguyen. Robust mapping of spatiotemporal trajectories and cell–cell interactions in healthy and diseased tissues. Nature communications, 14(1):7739–7739, 2023.

4. Suoqin Jin, Maksim V. Plikus, and Qing Nie. Cellchat for systematic analysis of cell-cell communication from single-cell and spatially resolved transcriptomics. bioRxiv, 2023.

5. Giovanni Palla, Hannah Spitzer, Michal Klein, David Fischer, Anna Christina Schaar, Louis Benedikt Kuemmerle, Sergei Rybakov, Ignacio L. Ibarra, Olle Holmberg, Isaac Virshup, Mohammad Lotfollahi, Sabrina Richter, and Fabian J. Theis. Squidpy: a scalable framework for spatial omics analysis. Nat Methods, 19(2):171–178, 2022.

6. Mirjana Efremova, Miquel Vento-Tormo, Sarah A. Teichmann, and Roser Vento-Tormo. Cellphonedb: inferring cell-cell communication from combined expression of multi-subunit ligand-receptor complexes. Nat Protoc, 15(4):1484–1506, 2020.

7. Rui Hou, Elena Denisenko, Huan Ting Ong, Jordan A. Ramilowski, and Alistair R. R. Forrest. Predicting cell-to-cell communication networks using natmi. Nature communications, 11(1):5011–5011, 2020.

8. Robin Browaeys, Wouter Saelens, and Yvan Saeys. Nichenet: modeling intercellular communication by linking ligands to target genes. Nat Methods, 17(2):159, 2020.

9. Cyril Lagger, Eugen Ursu, Anäis Equey, Roberto A. Avelar, Angela Oliveira Pisco, Robi Tacutu, and João Pedro de Magalhães. scdiffcom: a tool for differential analysis of cell-cell interactions provides a mouse atlas of aging changes in intercellular communication. Nat Aging, 3(11):1446– 1461, 2023.

10. Maria Solovey and Antonio Scialdone. Comunet: a tool to explore and visualize intercellular communication. Bioinformatics, 36(15):4296–4300, 2020.

11. Katy Vandereyken, Alejandro Sifrim, Bernard Thienpont, and Thierry Voet. Methods and applications for single-cell and spatial multi-omics. Nat Rev Genet, 24(8):494–515, 2023.

12. Shanshan He, Ruchir Bhatt, Carl Brown, Emily A. Brown, Derek L. Buhr, Kan Chantranuvatana, Patrick Danaher, Dwayne Dunaway, Ryan G. Garrison, Gary Geiss, Mark T. Gregory, Margaret L. Hoang, Rustem Khafizov, Emily E. Killingbeck, Dae Kim, Tae Kyung Kim, Youngmi Kim, Andrew Klock, Mithra Korukonda, Alecksandr Kutchma, Zachary R. Lewis, Yan Liang, Jeffrey S. Nelson, Giang T. Ong, Evan P. Perillo, Joseph C. Phan, Tien Phan-Everson, Erin Piazza, Tushar Rane, Zachary Reitz, Michael Rhodes, Alyssa Rosenbloom, David Ross, Hiromi Sato, Aster W. Wardhani, Corey A. Williams-Wietzikoski, Lidan Wu, and Joseph M. Beechem. High-plex multiomic analysis in ffpe at subcellular level by spatial molecular imaging. bioRxiv, page 2021.11.03.467020, 2022.

13. Amanda Janesick, Robert Shelansky, Andrew D. Gottscho, Florian Wagner, Stephen R. Williams, Morgane Rouault, Ghezal Beliakoff, Carolyn A. Morrison, Michelli F. Oliveira, Jordan T. Sicherman, Andrew Kohlway, Jawad Abousoud, Tingsheng Yu Drennon, Seayar H. Mohabbat, and Sarah E. B. Taylor. High resolution mapping of the tumor microenvironment using integrated single-cell, spatial and in situ analysis. Nat Commun, 14(1):8353–8353, 2023.

14. Kristen R. Maynard, Leonardo Collado-Torres, Lukas M. Weber, Cedric Uytingco, Brianna K. Barry, Stephen R. Williams, Joseph L. Catallini, Matthew N. Tran, Zachary Besich, Madhavi Tippani, Jennifer Chew, Yifeng Yin, Joel E. Kleinman, Thomas M. Hyde, Nikhil Rao, Stephanie C. Hicks, Keri Martinowich, and Andrew E. Jaffe. Transcriptome-scale spatial gene expression in the human dorsolateral prefrontal cortex. Nat Neurosci, 24(3):425– 436, 2021.

15. Ao Chen, Sha Liao, Mengnan Cheng, Kailong Ma, Liang Wu, Yiwei Lai, Xiaojie Qiu, Jin Yang, Jiangshan Xu, Shijie Hao, Xin Wang, Huifang Lu, Xi Chen, Xing Liu, Xin Huang, Zhao Li, Yan Hong, Yujia Jiang, Jian Peng, Shuai Liu, Mengzhe Shen, Chuanyu Liu, Quanshui Li, Yue Yuan, Xiaoyu Wei, Huiwen Zheng, Weimin Feng, Zhifeng Wang, Yang Liu, Zhaohui Wang, Yunzhi Yang, Haitao Xiang, Lei Han, Baoming Qin, Pengcheng Guo, Guangyao Lai, Pura Muñoz-Cánoves, Patrick H. Maxwell, Jean Paul Thiery, Qing-Feng Wu, Fuxiang Zhao, Bichao Chen, Mei Li, Xi Dai, Shuai Wang, Haoyan Kuang, Junhou Hui, Liqun Wang, Ji-Feng Fei, Ou Wang, Xiaofeng Wei, Haorong Lu, Bo Wang, Shiping Liu, Ying Gu, Ming Ni, Wenwei Zhang, Feng Mu, Ye Yin, Huanming Yang, Michael Lisby, Richard J. Cornall, Jan Mulder, Mathias Uhlén, Miguel A. Esteban, Yuxiang Li, Longqi Liu, Xun Xu, and Jian Wang. Spatiotemporal transcriptomic atlas of mouse organogenesis using dna nanoball-patterned arrays. Cell, 185(10):1777–1792.e21, 2022.

16. Pauli Virtanen, Ralf Gommers, Travis E. Oliphant, Matt Haberland, Tyler Reddy, David Cournapeau, Evgeni Burovski, Pearu Peterson, Warren Weckesser, Jonathan Bright, Stéfan J. van der Walt, Matthew Brett, Joshua Wilson, K. Jarrod Millman, Nikolay Mayorov, Andrew R. J. Nelson, Eric Jones, Robert Kern, Eric Larson, C. J. Carey, İlhan Polat, Yu Feng, Eric W. Moore, Jake VanderPlas, Denis Laxalde, Josef Perktold, Robert Cimrman, Ian Henriksen, E. A. Quintero, Charles R. Harris, Anne M. Archibald, Antônio H. Ribeiro, Fabian Pedregosa, and Paul van Mulbregt. Scipy 1.0: fundamental algorithms for scientific computing in python. Nat Methods, 17(3):261–272, 2020.

17. Shiyu Deng, Lin Gan, Chang Liu, Tongtong Xu, Shiyi Zhou, Yiyan Guo, Zhijun Zhang, Guo-Yuan Yang, Hengli Tian, and Yaohui Tang. Roles of ependymal cells in the physiology and pathology of the central nervous system. Aging Dis, 14(2):468–483, 2023.

18. Diana G. Nelles and Lili-Naz Hazrati. Ependymal cells and neurodegenerative disease: outcomes of compromised ependymal barrier function. Brain communications, 4(6):fcac288–fcac288, 2022.

19. Lilach Soreq, Jamie Rose, Eyal Soreq, John Hardy, Daniah Trabzuni, Mark R. Cookson, Colin Smith, Mina Ryten, Rickie Patani, and Jernej Ule. Major shifts in glial regional identity are a transcriptional hallmark of human brain aging. Cell Rep, 18(2):557–570, 2017.

20. Sergey V. Gudkov, Dmitriy E. Burmistrov, Elena V. Kondakova, Ruslan M. Sarimov, Roman S. Yarkov, Claudio Franceschi, and Maria V. Vedunova. An emerging role of astrocytes in aging/neuroinflammation and gut-brain axis with consequences on sleep and sleep disorders. Ageing Res Rev, 83:101775–101775, 2023.

21. Sreedevi Raman, Gayathri Srinivasan, Nicholas Brookhouser, Toan Nguyen, Tanner Henson, Daylin Morgan, Joshua Cutts, and David A. Brafman. A defined and scalable peptide-based platform for the generation of human pluripotent stem cell-derived astrocytes. ACS Biomater. Sci. Eng, 6(6):3477–3490, 2020.

22. Charles P. Couturier, Shamini Ayyadhury, Phuong U. Le, Javad Nadaf, Jean Monlong, Gabriele Riva, Redouane Allache, Salma Baig, Xiaohua Yan, Mathieu Bourgey, Changseok Lee, Yu Chang David Wang, V. Wee Yong, Marie-Christine Guiot, Hamed Najafabadi, Bratislav Misic, Jack Antel, Guillaume Bourque, Jiannis Ragoussis, and Kevin Petrecca. Single-cell rna-seq reveals that glioblastoma recapitulates a normal neurodevelopmental hierarchy. Nature communications, 11(1):1–19, 2020.

23. Natallia Makarava, Olga Mychko, Jennifer Chen-Yu Chang, Kara Molesworth, and Ilia V. Baskakov. The degree of astrocyte activation is predictive of the incubation time to prion disease. Acta Neuropathol Commun, 9(1):87–87, 2021.

24. Quan Yuan, Fen Wang, Sufang Xue, and Jianping Jia. Association of polymorphisms in the lrp1 and a2m genes with alzheimer’s disease in the northern chinese han population. Journal of Clinical Neuroscience, 20(2):253– 256, 2013.

25. Mitsuru Shinohara, Masaya Tachibana, Takahisa Kanekiyo, and Guojun Bu. Role of lrp1 in the pathogenesis of alzheimer’s disease: evidence from clinical and preclinical studies. Journal of lipid research, 58(7):1267–1281, 2017.

26. Petra May, Astrid Rohlmann, Hans H. Bock, Kai Zurhove, Jamey D. Marth, Eike D. Schomburg, Jeffrey L. Noebels, Uwe Beffert, J. David Sweatt, Edwin J. Weeber, and Joachim Herz. Neuronal lrp1 functionally associates with postsynaptic proteins and is required for normal motor function in mice. Mol Cell Biol, 24(20):8872–8883, 2004.

27. Hisako Muramatsu, Kun Zou, Nahoko Sakaguchi, Shinya Ikematsu, Sadatoshi Sakuma, and Takashi Muramatsu. Ldl receptor-related protein as a component of the midkine receptor. Biochem Biophys Res Commun, 270(3):936–941, 2000.

28. Mathew Tata, Christiana Ruhrberg, and Alessandro Fantin. Vascularisation of the central nervous system. Mechanisms of development, 138:26–36, 2015.

29. Christabel X. Tan and Cagla Eroglu. Cell adhesion molecules regulating astrocyte–neuron interactions. Curr Opin Neurobiol, 69:170–177, 2021.

30. Ulrike Beisiegel, Wilfried Weber, Gudrun Ihrke, Joachim Herz, and Keith K Stanley. The ldl–receptor–related protein, lrp, is an apolipoprotein e-binding protein. Nature, 341(6238):162–164, 1989.

31. Zhong Min Dai, Shuhui Sun, Chunyang Wang, Hao Huang, Xuemei Hu, Zunyi Zhang, Qing Richard Lu, and Mengsheng Qiu. Stage-specific regulation of oligodendrocyte development by wnt/β-catenin signaling. The Journal of neuroscience, 34(25):8467–8473, 2014.

32. Oligodendrocytes development and wnt signaling pathway. International journal of human anatomy.

33. Zsofia Miskolczi, Michael P. Smith, Emily J. Rowling, Jennifer Ferguson, Jorge Barriuso, and Claudia Wellbrock. Collagen abundance controls melanoma phenotypes through lineage-specific microenvironment sensing. Oncogene, 37(23):3166–3182, 2018.

34. Brett L. Ecker, Amanpreet Kaur, Stephen M. Douglass, Marie R. Webster, Filipe V. Almeida, Gloria E. Marino, Andrew J. Sinnamon, Madalyn G. Neuwirth, Gretchen M. Alicea, Abibatou Ndoye, Mitchell Fane, Xiaowei Xu, Myung Shin Sim, Gary B. Deutsch, Mark B. Faries, Giorgos C. Karakousis, and Ashani T. Weeraratna. Age-related changes in hapln1 increase lymphatic permeability and affect routes of melanoma metastasis. Cancer discovery, 9(1):82–95, 2019.

35. Cian D’Arcy and Christina Kiel. Cell adhesion molecules in normal skin and melanoma. Biomolecules (Basel, Switzerland), 11(8):1213, 2021.

36. Keiji Tanese, Yuuri Hashimoto, Zuzana Berkova, Yuling Wang, Felipe Samaniego, Jeffrey E. Lee, Suhendan Ekmekcioglu, and Elizabeth A. Grimm. Cell surface cd74–mif interactions drive melanoma survival in response to interferon-γ. Journal of investigative dermatology, 135(11):2775–2784, 2015.

37. Yan He, Miao Miao Sun, Guo Geng Zhang, Jing Yang, Kui Sheng Chen, Wen Wen Xu, and Bin Li. Targeting pi3k/akt signal transduction for cancer therapy. Signal transduction and targeted therapy, 6(1):425–425, 2021.

38. Fanglong Wu, Jin Yang, Junjiang Liu, Ye Wang, Jingtian Mu, Qingxiang Zeng, Shuzhi Deng, and Hongmei Zhou. Signaling pathways in cancer-associated fibroblasts and targeted therapy for cancer. Signal transduction and targeted therapy, 6(1):218–218, 2021.

39. Yun Yang, Wen-Long Ye, Ruo-Nan Zhang, Xiao-Shun He, Jing-Ru Wang, Yu-Xuan Liu, Yi Wang, Xue-Mei Yang, Yu-Juan Zhang, and Wen-Juan Gan. The role of tgf-β signaling pathways in cancer and its potential as a therapeutic target. Evidence-based complementary and alternative medicine, 2021:1–16, 2021.

40. Paola A Guerrero. Tgf-β activation and signaling in angiogenesis. In InTech eBooks, pages 3–23. IntechOpen, RIJEKA, 2017.

41. Peter M. Szabo, Amir Vajdi, Namit Kumar, Michael Y. Tolstorukov, Benjamin J. Chen, Robin Edwards, Keith L. Ligon, Scott D. Chasalow, Kin Hoe Chow, Aniket Shetty, Mohan Bolisetty, James L. Holloway, Ryan Golhar, Brian A. Kidd, Philip Ansumana Hull, Jeff Houser, Logan Vlach, Nathan O. Siemers, and Saurabh Saha. Cancer-associated fibroblasts are the main contributors to epithelial-to-mesenchymal signatures in the tumor microenvironment. Scientific reports, 13(1):3051–3051, 2023.

42. Dennis Pedri, Panagiotis Karras, Ewout Landeloos, Jean-Christophe Marine, and Florian Rambow. Epithelial-to-mesenchymal-like transition events in melanoma. The FEBS journal, 289(5):1352–1368, 2022.

43. Qinfeng Ma, Qiang Li, Xiao Zheng, and Jianbo Pan. Cellcommunet: an atlas of cell-cell communication networks from single-cell rna sequencing of human and mouse tissues in normal and disease states. Nucleic Acids Res, 52(D1), 2024.

44. Daniel Dimitrov, Dénes Türei, Martin Garrido-Rodriguez, Paul L. Burmedi, James S. Nagai, Charlotte Boys, Ricardo O. Ramirez Flores, Hyojin Kim, Bence Szalai, Ivan G. Costa, Alberto Valdeolivas, Aurélien Dugourd, and Julio Saez-Rodriguez. Comparison of methods and resources for cell-cell communication inference from single-cell rna-seq data. Nat Commun, 13(1):3224–3224, 2022.

45. Eleanor Catherine Sams. Oligodendrocytes in the aging brain. Neuronal signaling, 5(3):NS20210008–NS20210008, 2021.

46. Rosalía Fernández-Calle, Sabine C. Konings, Javier Frontiñań-Rubio, Juan García-Revilla, Lluís Camprubí-Ferrer, Martina Svensson, Isak Martinson, Antonio Boza-Serrano, José Luís Venero, Henrietta M. Nielsen, Gunnar K. Gouras, and Tomas Deierborg. Apoe in the bullseye of neurodegenerative diseases: impact of the apoe genotype in alzheimer’s disease pathology and brain diseases. Molecular neurodegeneration, 17(1):1–62, 2022.

47. Manuel Buttini, Eliezer Masliah, Gui-Qiu Yu, Jorge J. Palop, Shengjun Chang, Aubrey Bernardo, Carol Lin, Tony Wyss-Coray, Yadong Huang, and Lennart Mucke. Cellular source of apolipoprotein e4 determines neuronal susceptibility to excitotoxic injury in transgenic mice. Am J Pathol, 177(2):563–569, 2010.

48. Angela Marie Jablonski, Lee Warren, Marija Usenovic, Heather Zhou, Jonathan Sugam, Sophie Parmentier-Batteur, and Bhavya Voleti. Astrocytic expression of the alzheimer’s disease risk allele, apoeε4, potentiates neuronal tau pathology in multiple preclinical models. Sci Rep, 11(1):3438–18, 2021.

49. Natalie C. Schlegel, Anina von Planta, Daniel S. Widmer, Reinhard Dummer, and Gerhard Christofori. Pi3k signalling is required for a tgfβ-induced epithelial-mesenchymal-like transition (emt-like) in human melanoma cells. Experimental dermatology, 24(1):22–28, 2015.

50. Timothy Budden, Caroline Gaudy-Marqueste, Andrew Porter, Emily Kay, Shilpa Gurung, Charles H. Earnshaw, Katharina Roeck, Sarah Craig, Víctor Traves, Jean Krutmann, Patricia Muller, Luisa Motta, Sara Zanivan, Angeliki Malliri, Simon J. Furney, Eduardo Nagore, and Amaya Virós. Ultraviolet light-induced collagen degradation inhibits melanoma invasion. Nature communications, 12(1):2742–2742, 2021.

51. Léon C.L.T. van Kempen, Jos Rijntjes, Ine Mamor-Cornelissen, Silvia Vincent-Naulleau, Marie-Jeanne P. Gerritsen, Dirk J. Ruiter, Marcory C.R.F. van Dijk, Claudine Geffrotin, and Goos N.P. van Muijen. Type i collagen expression contributes to angiogenesis and the development of deeply invasive cutaneous melanoma. International journal of cancer, 122(5):1019–1029, 2008.

52. Shengchao Xu, Xizhe Li, Lu Tang, Zhixiong Liu, Kui Yang, and Quan Cheng. Cd74 correlated with malignancies and immune microenvironment in gliomas. Frontiers in molecular biosciences, 8:706949–706949, 2021.

53. Keiji Tanese, Yuuri Hashimoto, Zuzana Berkova, Yuling Wang, Felipe Samaniego, Jeffrey E. Lee, Suhendan Ekmekcioglu, and Elizabeth A. Grimm. Cell surface cd74–mif interactions drive melanoma survival in response to interferon-γ. J Invest Dermatol, 135(11):2775–2784, 2015.

54. Yasunari Fukuda, Matias A. Bustos, Sung-Nam Cho, Jason Roszik, Suyeon Ryu, Victor M. Lopez, Jared K. Burks, Jeffrey E. Lee, Elizabeth A. Grimm, Dave S. B. Hoon, and Suhendan Ekmekcioglu. Interplay between soluble cd74 and macrophage-migration inhibitory factor drives tumor growth and influences patient survival in melanoma. Cell death & disease, 13(2):117–117, 2022.

55. Dylan M. Cable, Evan Murray, Luli S. Zou, Aleksandrina Goeva, Evan Z. Macosko, Fei Chen, and Rafael A. Irizarry. Robust decomposition of cell type mixtures in spatial transcriptomics. Nat Biotechnol, 40(4):517–526, 2022.

56. Elaine H. Shen, Caroline C. Overly, and Allan R. Jones. The allen human brain atlas: Comprehensive gene expression mapping of the human brain. Trends Neurosci, 35(12):711– 714, 2012.

57. Yuhan Hao, Stephanie Hao, Erica Andersen-Nissen, William M. Mauck, Shiwei Zheng, Andrew Butler, Maddie J. Lee, Aaron J. Wilk, Charlotte Darby, Michael Zager, Paul Hoffman, Marlon Stoeckius, Efthymia Papalexi, Eleni P. Mimitou, Jaison Jain, Avi Srivastava, Tim Stuart, Lamar M. Fleming, Bertrand Yeung, Angela J. Rogers, Juliana M. McElrath, Catherine A. Blish, Raphael Gottardo, Peter Smibert, and Rahul Satija. Integrated analysis of multimodal single-cell data. Cell, 184(13):3573–3587.e29, 2021.

58. Ethel Shanas. The american soldier: Vol. i: Adjustment during army life. samuel a. stouffer the american soldier: Vol. ii: Combat and its aftermath. samuel a. stouffer. The American journal of sociology, 55(6):590–594, 1950.

59. Combining probability from independent tests: the weighted z-method is superior to fisher’s approach. Journal of evolutionary biology, 18(5):1368–1373, 2005.

60. F. Alexander Wolf, Philipp Angerer, and Fabian J. Theis. Scanpy: Large-scale single-cell gene expression data analysis. Genome Biology, 19(1):15–15, 2018.

61. Zhuoqing Fang, Xinyuan Liu, and Gary Peltz. Gseapy: a comprehensive package for performing gene set enrichment analysis in python. Bioinformatics, 39(1), 2023.

62. Aric A. Hagberg, Daniel A. Schult, and Pieter J. Swart. Exploring network structure, dynamics, and function using networkx. In Gäel Varoquaux, Travis Vaught, and Jarrod Millman, editors, Proceedings of the 7th Python in Science Conference, pages 11 – 15, Pasadena, CA USA, 2008.

63. Mihriban Karaayvaz, Simona Cristea, Shawn M. Gillespie, Anoop P. Patel, Ravindra Mylvaganam, Christina C. Luo, Michelle C. Specht, Bradley E. Bernstein, Franziska Michor, and Leif W. Ellisen. Unravelling subclonal heterogeneity and aggressive disease states in tnbc through single-cell rna-seq. Nature communications, 9(1):3588–10, 2018.

64. M. Claeson, S. X. Tan, D. Lambie, S. Brown, M. D. Walsh, P. D. Baade, N. Pandeya, K. J. Whitehead, H. P. Soyer, B. M. Smithers, D. C. Whiteman, and K. Khosrotehrani. The association between braf-v600e mutations and death from thin (≤1.00 mm) melanomas: A nested case–case study from queensland, australia. J Eur Acad Dermatol Venereol, 37(9):e1168–e1172, 2023.

65. Mihriban Karaayvaz, Simona Cristea, Shawn M. Gillespie, Anoop P. Patel, Ravindra Mylvaganam, Christina C. Luo, Michelle C. Specht, Bradley E. Bernstein, Franziska Michor, and Leif W. Ellisen. Unravelling subclonal heterogeneity and aggressive disease states in tnbc through single-cell rna-seq. Nat Commun, 9(1):3588–10, 2018.

